# Bispecific GD2xB7-H3 Antibody Improves Tumor Targeting and Reduces Toxicity while Maintaining Efficacy for Neuroblastoma

**DOI:** 10.64898/2026.09.21.753180

**Authors:** Amy K. Erbe, Arika S. Feils, Sabrina N. Reinthaler, Alina Hampton, Zachary T Rosenkrans, Yadira Guevara, Mildred Felder, Jessica Wiwczar, Daniel J. Gerhardt, Mark Bercher, Belinda Wenke, Callie Haertle, Mackenzie Heck, Kaia Heimstreet, Elizabeth Frankel, Megan Nielsen, Dan Spiegelman, Noah Tsarovsky, Jen Zaborek, Alexander L. Rakhmilevich, Jacquelyn A. Hank, Eduardo Aluicio-Sarduy, Jonathan W. Englev, Jonathan H. Davis, Bryan Glaser, Vladimir Subbotin, Roland Green, Reinier Hernandez, Bonnie Hammer, Paul M. Sondel

**Affiliations:** Dept. of Human Oncology, University of Wisconsin, Madison, WI, USA; Dept. of Medical Physics, University of Wisconsin, Madison, WI, USA; Invenra Inc., Madison WI, USA; Dept. of Biostatistics and Medical Informatics, Madison, WI, USA; Dept. of Pediatrics, University of Wisconsin, Madison, WI, USA

## Abstract

The current treatment regimen for neuroblastoma involves immunotherapy, including a monoclonal antibody that recognizes disialoganglioside (GD2), expressed at high levels on neuroblastoma. GD2 is not present on most normal tissues except for nerves. Thus, anti-GD2 antibody treatment causes substantial, dose-limiting neuropathic pain. B7-H3 is overexpressed on multiple tumor types, including neuroblastoma, and is largely absent on nerves and other normal tissues. We designed a bispecific antibody (bsAb) that requires simultaneous binding of these two tumor antigens to achieve tight binding of tumor cells. Our preclinical research shows that, compared with a monospecific anti-GD2 antibody, the GD2×B7-H3 bsAb has improved tumor specificity, comparable antitumor efficacy, and reduced nerve binding and pain-associated toxicity. Since this bsAb does not bind to nerves, it may permit more tolerable and sustained treatment schedules than are currently feasible with monospecific anti-GD2 antibodies. In addition, its enhanced tumor specificity may support future development as a targeted delivery platform for antibody-drug conjugates or other payload-based therapies, potentially improving both efficacy and quality of life for patients with neuroblastoma.

## INTRODUCTION

Neuroblastoma (NBL) is the most common extracranial solid tumor in children, affecting approximately 800 children annually in the United States, nearly half of which present with high-risk disease. Despite intensive multimodal therapy, high-risk neuroblastoma (HR-NBL) retains a poor prognosis, with a survival rate of ∼60%. Current treatment consists of three phases: i) induction therapy including multi-agent chemotherapy and surgery; ii) consolidation using myeloablative chemotherapy, autologous stem cell transplantation and radiation therapy; and iii) maintenance therapy aimed at eradicating residual disease with immunotherapy.^1^ Maintenance immunotherapy includes an anti-GD2 monoclonal antibody (mAb), most commonly dinutuximab, administered with granulocyte-macrophage colony-stimulating factor (GM-CSF) and retinoic acid, and has significantly improved event-free and overall survival in children with HR-NBL.^1, 2^

The antitumor activity of dinutuximab is mediated primarily through antibody-dependent cellular cytotoxicity (ADCC), involving natural killer (NK) cells, neutrophils, and monocytes/macrophages.^3, 4^ Clinical studies have demonstrated that anti-GD2 mAbs, including dinutuximab and related agents, such as naxitamab, dinutuximab-beta, and hu14.18K322A, provide meaningful therapeutic benefit in NBL.^5, 6, 7, 8^ However, clinical use of anti-GD2 therapy is limited by severe neuropathic pain experienced by nearly all patients due to GD2 expression on peripheral nerves.^2, 9, 10^ Indeed, this neuropathic toxicity was the dose-limiting adverse event that established the maximum tolerated dose of dinutuximab in patients. Consequently, anti-GD2 mAbs are administered at substantially lower doses than other therapeutic mAbs, including rituximab and cetuximab.^11, 12^ These dose-limiting toxicities highlight a major challenge in NBL immunotherapy: improving the therapeutic index of GD2-directed therapy while minimizing on-target, off-tumor toxicity resulting from GD2 expression on normal tissues.

Importantly, several clinical studies have reported a positive association between anti-GD2 antibody exposure and patient outcomes, suggesting that strategies capable of improving tolerability may facilitate more sustained therapeutic exposure.^6, 13, 14^ In addition, highly tumor-selective targeting approaches could provide a foundation for future payload-based applications, including antibody-drug conjugates (ADCs) and radiopharmaceuticals. To overcome the dose-limiting neuropathic toxicity associated with conventional anti-GD2 therapy, we developed a human IgG bispecific antibody (bsAb), GD2xB7-H3, targeting both GD2 and B7-H3 (CD276), a surface protein highly expressed on NBL but minimally expressed on normal tissues and absent from peripheral nerves.^15, 16^ In this study, we evaluated the antigen specificity, tumor-targeting capacity, immune effector function, antitumor efficacy, and pain-associated toxicity of the fucosylated (INV721) and afucosylated (INV724) forms of this GD2xB7-H3 bsAb in a series of *in vitro* and *in vivo* NBL models. Our findings demonstrate that dual-antigen targeting preserves potent antitumor activity while augmenting the antitumor specificity, thereby substantially reducing nerve binding and neuropathic toxicity, supporting the clinical development of this strategy for NBL immunotherapy.

## RESULTS

### *In Vitro* Antigen/Target Specificity

To enhance anti-GD2 tumor specificity, we leveraged Invenra’s B-Body® platform to engineer a human IgG containing one Fragment antigen-binding (Fab) arm recognizing GD2 and a second Fab arm recognizing B7-H3 (**Fig. 1a and Fig. S1a**). Multiple anti-GD2 and anti-B7-H3 Fab combinations were screened to identify a format that achieved selective binding of cells co-expressing both antigens while minimizing binding to cells expressing either antigen alone. The optimized GD2xB7-H3 bsAb incorporates a low-affinity anti-GD2 arm (monovalent *K*d >1 μM) and a moderate-affinity anti-B7-H3 arm (*K*d = 2 nM). Because each Fab arm of the GD2xB7-H3 bsAb possesses only weak-to-moderate monomeric affinity, the antibody binds inefficiently to cells expressing either target alone (i.e. GD2**^+^**/B7-H3**^-^** peripheral nerves or GD2**^-^**/B7-H3**^+^** liver cells). Instead, stable cell-surface binding is driven by avidity and requires co-expression of both GD2 and B7-H3 (**Fig. 1b-e**).^175^ In contrast, dinutuximab binds GD2 with high affinity (*K*d = 10 nM)^18^, enabling efficient recognition of GD2-expressing cells regardless of B7-H3 expression, including peripheral nerves (**Fig. 1a**).

**Figure 1.**
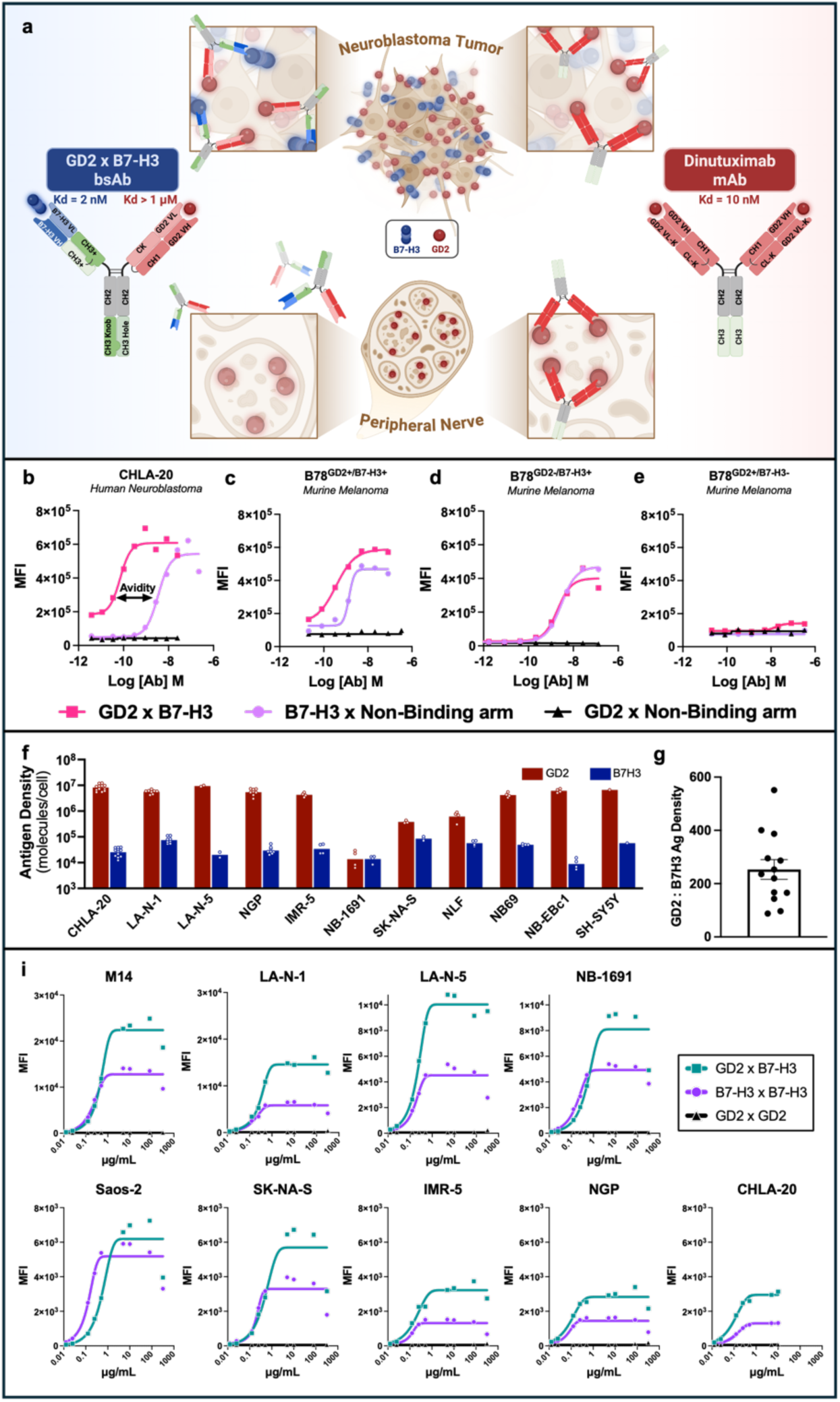
GD2xB7-H3 bsAb exhibits dual-antigen–dependent high-avidity binding to NBL in vitro. **a**) Dinutuximab binds GD2 with high affinity (Kd = 10 nM) and recognizes both GD2⁺ tumors and GD2⁺ nerves through bivalent GD2 engagement. In contrast, the GD2×B7-H3 bsAb (INV721) contains a low-affinity anti-GD2 Fab (Kd > 1 μM) and a moderate-affinity anti-B7-H3 Fab (Kd = 2 nM), resulting in high-avidity binding only to GD2⁺B7-H3⁺ NBL cells and not to GD2⁺B7-H3⁻ nerves. **b-e**) Flow cytometry comparing GD2×B7-H3 bsAb (INV721) with monospecific controls (B7-H3 Fab × non-binding Fab; GD2 Fab × non-binding Fab). The GD2×B7-H3 bsAb demonstrated enhanced binding to human CHLA-20 neuroblastoma cells relative to monospecific antibodies (**b**). B78 melanoma variants showed enhanced binding of GD2×B7-H3 bsAb to GD2⁺B7-H3⁺ cells (**e**), binding comparable to the B7-H3 Fab × non-binding Fab control on GD2⁻B7-H3⁺ cells (**d**), and minimal binding to GD2⁺B7-H3⁻ cells (**e**). **f-g)** Antigen expression analysis of 11 human NBL cell lines demonstrated co-expression of GD2 and B7-H3, with GD2 expressed at approximately 200-fold higher antigen density than B7-H3 (**f**). A dot plot shows the ratio of antigen density for GD2-to-B7-H3, with dot representing one of the 11 NBL cell lines shown in **f** (**g**). **h**) Antibody titration studies comparing GD2×B7-H3 bsAb (INV724), I7-01 mAb (B7-H3xB7-H3 mAb), and GD2-7 mAb (GD2xGD2 mAb) across 9 human pediatric tumor cell lines (7 NBL, M14 melanoma, and Saos-2 osteogenic sarcoma).

To distinguish the affinity of the individual Fab arms from the avidity of dual-antigen recognition, we generated single-arm control antibodies consisting of: (i) the B7-H3 Fab paired with a non-binding Fab arm (B7-H3xNon-Binding) or (ii) the GD2 Fab paired with a non-binding Fab arm (GD2xNon-Binding). We then compared the binding characteristics of these control antibodies with those of the GD2xB7-H3 bsAb using CHLA-20 human NBL (endogenously GD2**^+^**/B7-H3**^+^**) and B78 murine melanoma cells engineered to express GD2 and/or human B7-H3 (B78^GD2+/B7-H3+^, B78^GD2+/B7-H3-^, and B78^GD2-/B7-H3+^). On CHLA-20 and B78^GD2+/B7-H3+^, GD2xB7-H3 bsAb demonstrated robust binding at low concentrations (**Fig. 1b-c**), consistent with avidity-mediated dual-antigen recognition. In contrast, B7-H3xNon-Binding antibody showed detectable binding only at higher concentrations, reflecting the moderate monovalent affinity of the anti-B7-H3 Fab. The GD2xNon-Binding antibody showed no detectable binding, even at the highest concentrations tested, demonstrating that the monovalent anti-GD2 arm alone is insufficient to mediate appreciable cell-surface binding. On B78^GD2-/B7-H3+^ cells, both GD2xB7-H3 and B7-H3xNon-Binding antibodies exhibited detectable binding only at higher concentrations, whereas the GD2xNon-Binding antibody showed no binding (**Fig. 1d**). Conversely, on B78^GD2+/B7-H3-^cells none of these three antibodies demonstrated detectable binding, even at high concentrations (**Fig. 1e**). Together, these findings demonstrate that stable, high-avidity binding by the GD2xB7-H3 bsAb is dependent on simultaneous recognition of GD2 and B7-H3 and cannot be achieved through either monovalent Fab arm alone.

As reported previously by Majzner et al., GD2 and B7-H3 are co-expressed across most human NBL cell lines.^19^ We tested 11 HR-NBL cell lines and observed variable levels of expression of GD2 and B7-H3 across the panel (**Fig. 1f**). In most lines, GD2 was expressed at substantially higher antigen density than B7-H3 (**Fig. 1f**), with an average ∼200-fold greater expression of GD2 across all cell lines tested (**Fig. 1g**). Despite this variability, the GD2xB7-H3 bsAb bound all HR-NBL cell lines evaluated (**Fig. 1h**). Since there is roughly 200-fold more GD2 than B7-H3 on most human NBL cell lines (**Fig. 1g**), at saturating conditions, the amount of dinutuximab that can bind to these human NBL cells is greater than the amount of GD2xB7-H3 bsAb that can bind (**Fig. S1b-d**). To further assess the contribution of each Fab arm of GD2xB7-H3 bsAb, we generated two monospecific Abs from the bsAb format containing either two anti-GD2 Fab arms (GD2-7) or two anti-B7-H3 Fab arms (I7-01). Although binding avidity varied among 7 HR-NBL cell lines and a human melanoma and osteosarcoma line, GD2xB7-H3 consistently demonstrated stronger binding at lower concentrations, and greater maximal binding than I7-01 (**Fig. 1h**). This enhanced binding may reflect the differential ratio of GD2 and B7-H3 antigen density on the cell surface (**Fig. S1e).** Of note, minimal binding by GD2-7 was observed across these 9 cell lines, consistent with the intentionally low affinity of the anti-GD2 Fab incorporated into the GD2-B7H3 bsAb (**Fig. 1h**).

### *In Vitro* Immune-Mediated Antitumor Efficacy

A primary mechanism of anti-GD2 mAb activity is the induction of ADCC against GD2^+^ cells.^20, 21^ ADCC is mediated by Fc gamma receptor (Fc*γ*R)-bearing immune cells that recognize the Fc region of tumor-bound antibodies and subsequently induce cytotoxicity. Previous studies have reported that anti-GD2 mAbs, including dinutuximab, internalize within 16-24 hours after binding to target cells, which may represent a mechanism of immune resistance.^22^ Consequently, antibodies that exhibit limited or delayed internalization may enhance ADCC by maintaining cell-surface Fc exposure and facilitating sustained engagement of immune effector cells. Consistent with these findings, we observed internalization of dinutuximab over 16 hours in five human tumor cell lines that endogenously express GD2 and B7-H3 (**Fig. 2a**). In contrast, minimal internalization was observed for GD2xB7-H3 bsAb or anti-B7-H3 mAb (I7-01) in this same assay (**Fig. 2b**). Despite internalization, dinutuximab retained potent ADCC activity at 50 ng/mL, with efficacy comparable to that of GD2xB7H3, which contains the same human IgG1 Fc backbone as dinutuximab (**Fig. 2b**).

**Figure 2.**
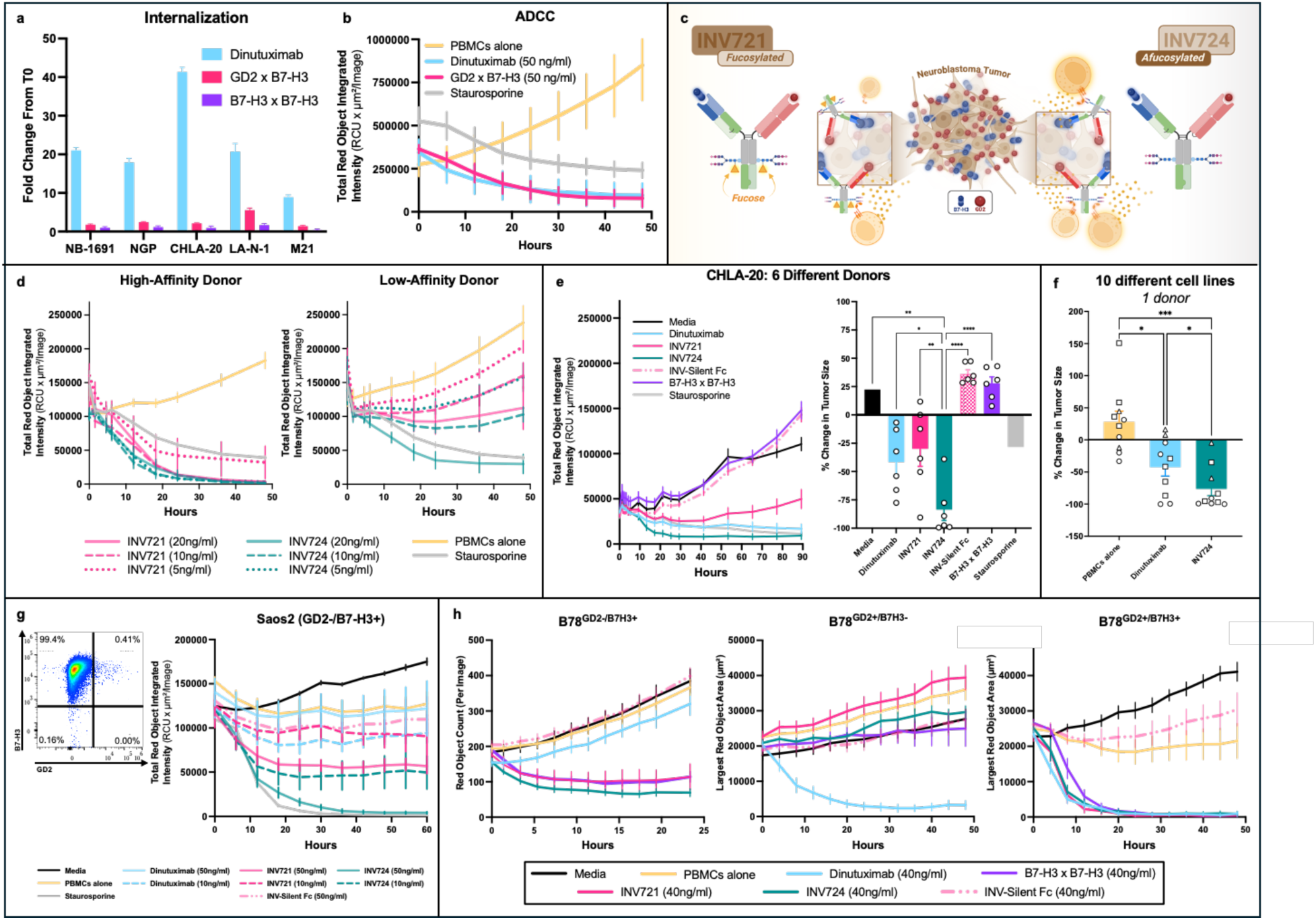
GD2 × B7-H3 bispecific antibodies efficiently mediate ADCC with reduced off-target activity. **a**) Across five human tumor cell lines, dinutuximab internalizes more extensively than GD2xB7-H3 bsAb (INV721) or anti–B7-H3 mAb (I7-01), quantified as fold change of internalized mAb over baseline. **b**) Using a high FcγR affinity donor, ADCC assays via IncuCyte show that GD2xB7-H3 bsAb (INV721) and dinutuximab induce comparable spheroid shrinkage of CHLA-20 neuroblastoma cells by PBMCs, while PBMCs alone do not mediate cytotoxicity. **c**) GD2×B7-H3 bsAb was generated with two distinct Fc glycosylation profiles: INV721, which contains a conventionally fucosylated Fc domain, and INV724, with an afucosylated Fc domain. **d**) ADCC potency against the CHLA-20 NBL depends on FcγR genotype and Fc fucosylation: INV724 (afucosylated Fc) mediates robust killing across concentrations for both donors whereas INV721 (fucosylated Fc) shows reduced ADCC activity particularly with low-affinity FcγR PBMCs. **e**) PBMCs from six different donors showed GD2×B7-H3 bsAbs (INV721, INV724) and dinutuximab induce significant ADCC at 50 ng/ml against CHLA-20, whereas an effector-silent Fc control does not; tumor growth (left plot) and % change in tumor size (right plot). **f)** Using effector cells from a single PBMC donor with antibody concentrations at 50 ng/ml, INV724 elicited significantly greater ADCC compared to dinutuximab after 24 hrs using 11 different tumor cell lines, including 4 melanomas (circles), 4 NBL (triangles), and 2 sarcomas (Ewing sarcoma and osteosarcoma; squares). **g**) In GD2⁻B7-H3⁺ Saos-2 osteosarcoma cells, INV724, but not INV721 or dinutuximab, induced potent ADCC at 50 ng/ml; efficacy of ADCC was reduced with 10 ng/ml INV724. **h**) INV724 is capable of eliciting ADCC against murine B78 melanoma cells that express human B7H3+ but lack GD2 expression (left), INV724 and dinutuximab mediate ADCC of murine B78 melanoma cells engineered to express GD2 and human B7-H3 (right), whereas B78 cells expressing only GD2 but not B7-H3 (middle) are killed by dinutuximab but not INV724, demonstrating tumor-restricted cytotoxicity. ****p<0.0001; **p<0.01; *p<0.05.

The interaction between FcγRs and tumor-bound antibodies is strongly influenced by Fc fucosylation, whereby reduced fucosylation enhances FcγR binding by 10-100 fold.^23^ To investigate the impact of Fc fucosylation on GD2xB7-H3-mediated engagement, we generated two variants of the GD2xB7-H3 bsAb with identical Fab binding domains but different Fc fucosylation profiles (**Fig. 2c**). The fucosylated variant, INV721, contains a conventional IgG1 Fc region reflecting that of dinutuximab, whereas the afucosylated INV724 was designed to augment binding to FcγRs on NK cells and enhance ADCC-mediated tumor killing (**Fig. 2c**).

We and others have shown that response to mAb immunotherapy can vary based on an individual’s FcγR genotype.^24, 25, 26^ Using NK cells from a donor with a high-affinity FcγR genotype, we found that INV721 and INV724 elicited similar ADCC at high antibody concentrations (20 and 10 ng/mL). However, at a low concentration (5 ng/mL), INV721 exhibited less ADCC than INV724 (**Fig. 2d**, left; **Fig. S2A**). In contrast, when NK cells from a donor with a low-affinity FcγR genotype were used, INV724 retained ADCC activity at 20 ng/mL, whereas INV721 largely lost its potency (**Fig. 2d**, right). Moreover, when evaluated across NK cells from six independent donors, INV724 mediated significantly greater ADCC against CHLA-20 NBL cells than either dinutuximab or INV721 (**Fig. 2e**). Further, INV724 mediated greater ADCC than dinutuximab in 10 different tumor cell lines (4 NBL, 4 melanoma, 1 osteosarcoma, and 1 Ewing sarcoma) (**Fig. 2f, Fig. S2b**). Based on these findings, INV724 was selected as the lead candidate and used for subsequent *in vivo* efficacy studies.

Although the avidity of INV724 enables potent ADCC against GD2^+^/B7-H3^+^ tumor cells, the moderate monovalent affinity of its anti-B7-H3 Fab also permits binding to GD2^-^/B7-H3^+^ cells at higher antibody concentrations (**Fig. 1c-e**). Consistent with monovalent B7-H3-mediated activity at higher concentrations, INV724 mediated robust ADCC against GD2^-^/B7-H3^+^ Saos-2 cells at 50 ng/mL, but substantially less activity at 10 ng/mL (**Fig. 2g**). INV724 also induced greater ADCC against B78^GD2**-**/B7-H3**+**^ cells at 50 ng/mL than either INV721 or I7-01 (anti-B7-H3 mAb), both of which contain fucosylated Fc domains (**Fig. 2h**, left). In contrast, the low-affinity anti-GD2 Fab arm of INV724 does not support binding to GD2^+^/B7-H3^-^cells. B78^GD2**+**/B7-H3**-**^cells, which mimic the GD2^+^/B7-H3^-^phenotype of peripheral nerve cells, were susceptible to ADCC mediated by dinutuximab but not INV724 (**Fig. 2h**, middle), whereas both dinutuximab and INV724 mediated efficient killing of B78^GD2**+**/B7-H3**+**^ cells (**Fig. 2h**, right). Similar patterns of ADCC were observed at lower antibody concentrations, and B78^GD2**-**/B7-H3**-**^cells were not susceptible to ADCC mediated by either antibody (**Fig. S2c**). Thus, despite equivalent GD2 expression on the GD2^+^/B7-H3^+^ and GD2^+^/B7-H3^-^B78 variants, INV724 mediated ADCC only when B7-H3 was co-expressed, demonstrating that dual-antigen recognition effectively restricts cytotoxicity toward GD2^+^/B7-H3^+^ target cells.

### INV721 *In Vivo* Tumor-Targeting and Specificity

To test *in vivo* tumor targeting, we assessed *in vivo* trafficking of GD2xB7-H3 using positron emission tomography (PET) imaging of Zirconium-89 (^89^Zr) radiolabeled mAbs injected intravenously (IV) in NRG mice bearing human NGP neuroblastoma xenografts (which endogenously express GD2 and B7-H3; **Fig. 1f**). We found that ^89^Zr-INV721 (fucosylated GD2×B7-H3) localized to and was retained in established NGP tumors, with significantly more uptake than ^89^Zr-dinutuximab (**Fig. 3a-b; Fig. S3a**). Minimal uptake was found in other tissue, except for uptake within the spleen for both control (^89^Zr-B-Body) and ^89^Zr-INV721 (**Fig. S3b**), consistent with nonspecific clearance of radiolabeled antibodies by the mononuclear phagocyte system.^27^

**Figure 3.**
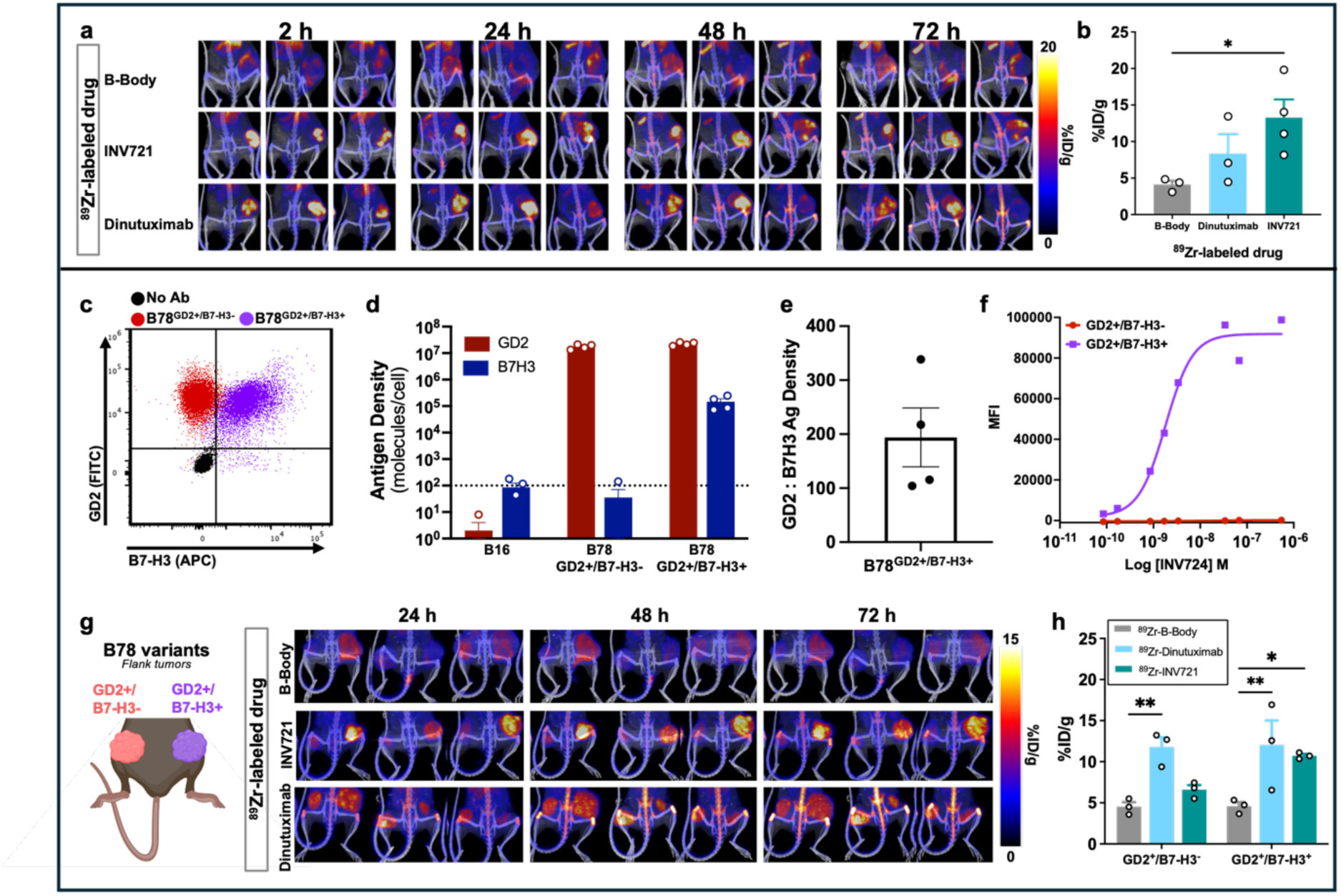
INV721 selectively targets GD2⁺B7-H3⁺ tumors with high avidity in vivo. **a)** Immuno-PET imaging demonstrates tumor uptake and prolonged retention of ^89^Zr-INV721 in NGP xenografts, with greater accumulation than ^89^Zr-dinutuximab and no tumor uptake observed for a non-binding control antibody. ^89^Zr-BBody serves as a negative antibody control. **b)** Quantitative PET analysis demonstrates significantly greater tumor uptake of ^89^Zr-INV721 compared with ^89^Zr-dinutuximab in NGP xenografts. **c)** Flow cytometric contour plots showing GD2 and B7-H3 surface expression on B78^GD2⁺B7-H3⁺^ and B78^GD2⁺B7-H3⁻^ melanoma variants. **d)** Antigen density analysis (log scale) demonstrates that B78^GD2⁺B7-H3⁺^ cells express GD2 at a higher density than B7-H3, GD2 is expressed at a comparable density for both B78^GD2⁺B7-H3-^ and B78^GD2⁺B7-H3⁺^ cells. B16 melanoma cells lacking GD2 and human B7-H3 expression served as a negative control. **e)** The GD2:B7-H3 antigen density ratio in B78^GD2⁺B7-H3⁺^ cells (n=4) is approximately 200:1, comparable to that observed in human NBL cell lines (Fig. 1g). **f)** In vitro titration demonstrates selective binding of GD2×B7-H3 bsAb INV724 to B78^GD2⁺B7-H3⁺^ cells, with minimal binding to B78^GD2⁺B7-H3-^cells. **g)** Schematic of bilateral tumor implantation showing B78^GD2⁺B7-H3⁺^ tumors on the right flank and B78^GD2⁺B7-H3⁻^ tumors on the left flank. Immuno-PET imaging demonstrates preferential uptake of ^89^Zr-INV721 in B78^GD2⁺B7-H3⁺^ tumors, on the right at 24, 48 and 72h whereas ^89^Zr-dinutuximab accumulates in both B78^GD2⁺B7-H3⁺^ and B78^GD2⁺B7-H3⁻^ tumors at these same times. **h)** Quantitative biodistribution analysis demonstrates significantly greater accumulation of ^89^Zr-INV721 in B78^GD2⁺B7-H3⁺^ tumors than ^89^Zr-dinutuximab.

To assess the *in vivo* specificity of this trafficking, we evaluated ^89^Zr-INV721 uptake into tumor variants with differential GD2 and B7-H3 expression. Using B78^GD2+/B7-H3-^ and B78^GD2+/B7-H3+^ cell lines, which both express GD2 but only the latter expresses B7-H3 (**Fig. 3c**), we quantified GD2 and B7-H3 antigen density on each variant (**Fig. 3d**) and confirmed that B78^GD2+/B7-H3+^ cells exhibited a GD2:B7-H3 ratio (**Fig. 3e**) comparable to that observed in HR-NBL cell lines (**Fig. 1g**). Consistent with its bispecific design, *in vitro* titration demonstrated that GD2xB7-H3 bsAb bound selectively to the B78^GD2+/B7-H3+^ cells, with no detectable binding to B78^GD2+/B7-H3-^cells (**Fig. 3f**). To evaluate antigen-dependent targeting *in vivo*, mice were implanted with B78^GD2+/B7-H3-^tumors on the left flank and B78^GD2+/B7-H3+^ on the right flank (**Fig. 3g**). PET imaging revealed selective accumulation of ^89^Zr-INV721 in B78^GD2+/B7-H3+^ tumors, whereas ^89^Zr-dinutuximab accumulated in both tumor types (**Fig. 3g-h**), demonstrating that tumor localization of GD2xB7-H3 bsAb requires co-expression of GD2 and B7-H3. Complete biodistribution (BioD) data are shown in **Fig. S3c**. Together, these findings demonstrate that INV721 retains efficient trafficking to dual-antigen tumors while discriminating against GD2-positive tissues lacking B7-H3, supporting its potential to reduce on-target, off-tumor toxicities associated with GD2-directed therapies.

### *In Vivo* Antitumor Immunotherapeutic Efficacy

Using syngeneic murine models for both NBL and melanoma, we observed robust *in vivo* antitumor efficacy of INV724 in mice bearing GD2^+^/B7-H3^+^ tumors. GD2-expressing murine tumor models, including B78 melanoma (syngeneic to C57BL/6 mice) and NXS2 neuroblastoma (syngeneic to A/J mice), were transduced to express human B7-H3 and treated using an in situ vaccine regimen developed in our laboratory consisting of radiation therapy (RT), tumor-targeting antibody, and the immunostimulatory cytokine IL-2.^28, 29^ In mice bearing B78^GD2+/B7-H3+^ tumors (**Fig. 4a-c)** or mice bearing NXS2 ^GD2+/B7-H3+^ tumors (**Fig. 4d-f**), the addition of INV724 to RT + IL-2 significantly improved antitumor responses compared with RT + IL-2 alone (**Fig. 4b-c and 4e-f**).

**Figure 4.**
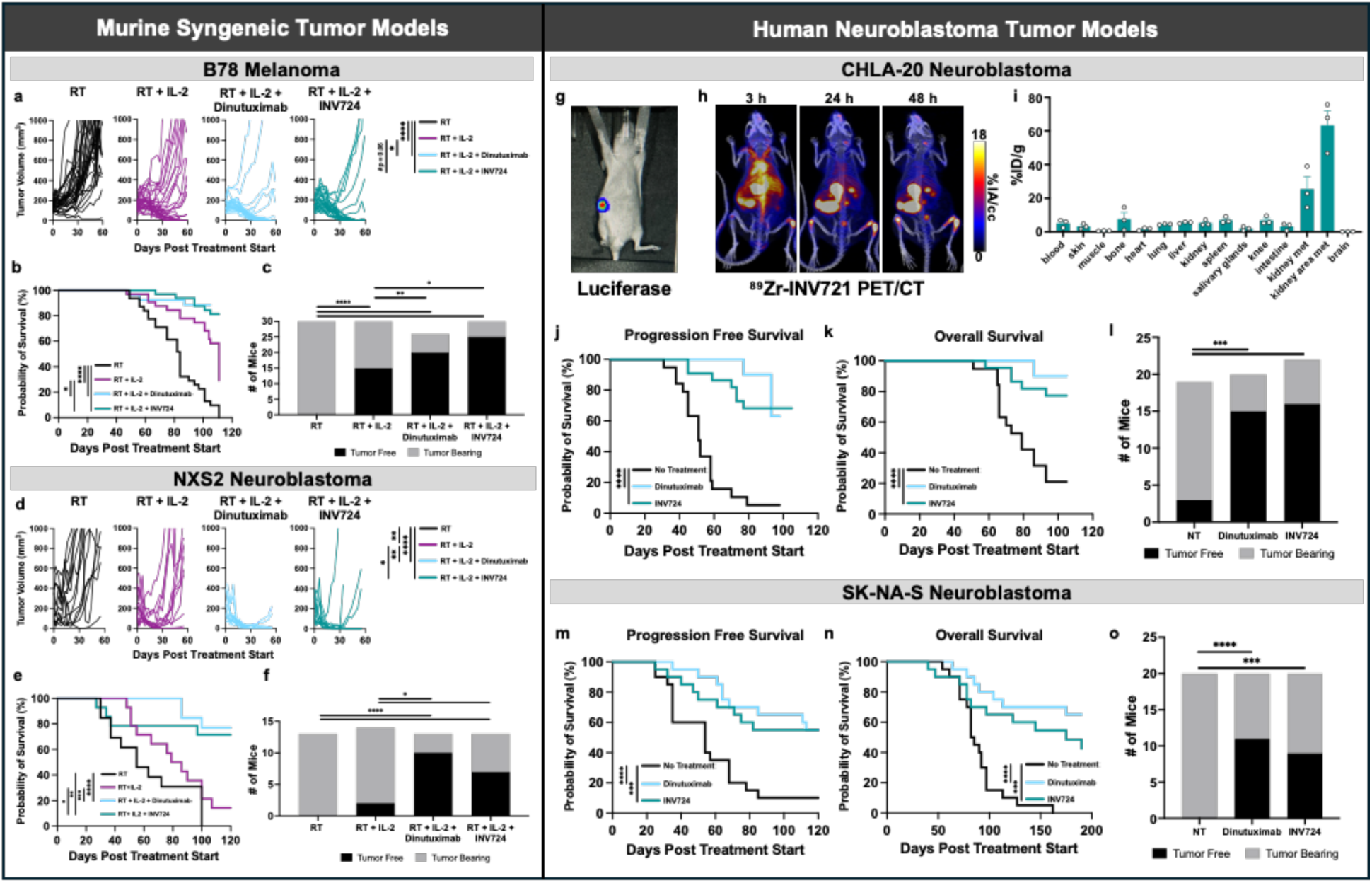
INV724 mediates antitumor efficacy in murine solid tumor models. **a-f)** In syngeneic B78 melanoma (**a-c**) and NXS2 neuroblastoma (**d-f**) models, mice treated with RT plus INV724 or dinutuximab in combination with IL-2 show reduced tumor growth (**a-d**), improved overall survival (**b-e**), and increased tumor-free mice (**c-f**) compared with RT alone or RT + IL-2. **g-l)** In the IV-inoculated metastatic human NBL CHLA-20-luc xenograft model, CHLA-20-luc tumors preferentially localized to a kidney based on luciferase imaging (**g**), and PET imaging showed accumulation of ⁸⁹Zr-INV724 within sites of metastatic disease (**h**) confirmed by quantitative BioD (**i**). Treatment with either INV724 or dinutuximab significantly delayed disease progression as assessed by IVIS imaging resulting in improved PFS **(j)** and improved OS **(k)** relative to the NT control. INV724 and dinutuximab also increased the proportion of tumor-free mice (**l**). **m-o**) Using a metastatic human neuroblastoma SK-NA-S-luc xenograft model, treatment with INV724 or dinutuximab similarly resulted in significantly improved PFS (**m**), OS (**n**), and increased numbers of tumor-free mice (**o**) compared with untreated controls. For **a-f** and **j-n** data represent pooled results from two independent experiments per model. ****p<0.0001; ***p<0.001; **p<0.01; *p<0.05.

We next evaluated the single-agent activity of INV724 in nude mice bearing human NBL experimental-metastases established by intravenous (IV) inoculation of luciferase-expressing tumor cells. Treatment with INV724 significantly improved progression-free survival (PFS), overall survival (OS), and the proportion of tumor-free mice compared with untreated controls in two separate tumor models (**Fig. 4g-o**). IV-injected CHLA-20 tumor cells established disease predominantly within the abdominal cavity, including the kidneys (**Fig. 4g**), and ^89^Zr-INV724 trafficked to these disease sites with minimal off-tumor uptake observed (**Fig. 4h-i**). Treatment of CHLA-20 (**Fig. 4j-l)** or SK-NA-S (**Fig. 4m-o**) tumor-bearing mice with single-agent antibody therapy (dinutuximab or INV724) significantly improved PFS, OS, and the proportion of tumor-free mice compared to untreated controls. No statistically significant differences were observed between the efficacy of INV724 and dinutuximab in any of these studies (**Fig. 4**). Together, these *in vivo* tumor-targeting (**Fig. 3**) and antitumor efficacy studies (**Fig. 4**) demonstrate the potential utility of INV724 as an alternative to dinutuximab for cancer imaging and immunotherapy.

### *Ex Vivo* Nerve Binding and *In Vivo* Pain Toxicity Studies

In addition to tumors, GD2 is also expressed on human central nervous system cells and the myelin sheaths of peripheral nerves.^30, 31^ GD2 expression on peripheral nerves is implicated in the peripheral neuropathic pain experienced by patients during dinutuximab therapy.^32^ To determine whether the GD2xB7-H3 bsAbs (INV721 and INV724), which contain identical antigen-binding domains, are incapable of binding to nerve tissue, we performed immunofluorescent staining of neural tissues. Dinutuximab exhibited detectable binding to rat sympathetic ganglion neurons, whereas INV721 showed no detectable binding (**Fig. 5a**). Similarly, dinutuximab also bound human peripheral nerve fibers, while INV721 did not (**Fig. 5b**). In contrast, both dinutuximab and INV721 demonstrated comparable binding to CHLA-20 neuroblastoma xenograft tissue (**Fig. 5a-b**).

**Figure 5.**
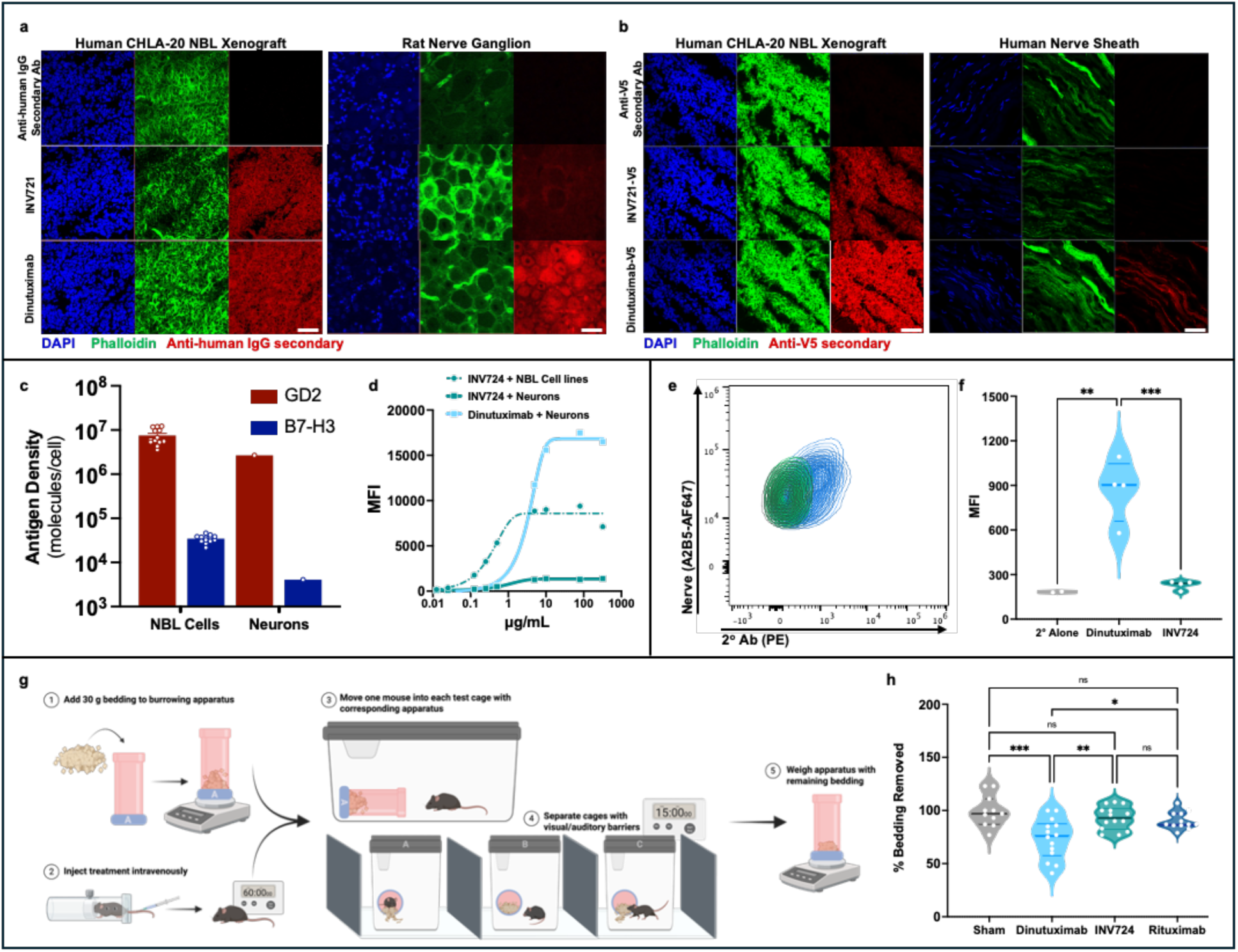
Anti-GD2×B7-H3 bispecific antibody does not bind peripheral nerves and does not induce pain behavior in mice, whereas dinutuximab does. **a-b)** Immunofluorescence staining of CHLA-20 xenograft tumor and peripheral rat sympathetic ganglion (**a**) and CHLA-20 xenograft tumor and human peripheral nerves (**b**), using DAPI (blue), phalloidin (green), and either INV721, dinutuximab, or control antibody, followed by detection of human anti-IgG (red; **a**) or anti-V5 tag (red; **b**). Both antibodies bound CHLA-20 tumors; however, only dinutuximab bound rat sympathetic ganglia and human peripheral nerves, whereas INV721 showed no detectable nerve binding. **c-d**) Flow cytometric analysis of human neurons demonstrating GD2 expression at lower antigen density than for HR-NBL cells (n=11 separate NBL lines; Fig. 1f) and minimal B7-H3 expression (**c**). Antibody titration studies showed potent binding of dinutuximab to neurons, and of INV724 to human NBL cells (n=8 NBL lines, Fig. S1) whereas INV724 exhibited minimal neuronal binding across the concentration range tested (**d**). **e-f**) Flow cytometric analysis of cells isolated from murine dorsal root ganglia (n = 4 ganglia/group) labeled with anti-ganglioside (A2B5) demonstrated substantial binding of dinutuximab but minimal binding of INV724, as detected by fluorescent anti-human IgG secondary Ab (**e**), with significantly greater MFI for dinutuximab compared to INV724 and secondary Ab alone (**f**). **g-h**) Burrowing assay in C57BL/6 mice following intravenous administration of INV724, dinutuximab, rituximab, or sham control. Burrowing behavior was assessed 1 h after treatment relative to baseline. Burrowing activity was quantified as the percentage of bedding displaced during the assay period (**g**). INV724-treated mice exhibited burrowing activity comparable to rituximab-treated and sham-treated controls, whereas dinutuximab-treated mice exhibited significantly reduced burrowing behavior than all 3 other groups, consistent with pain-associated suppression of normal activity (**h**). Data in h represent three independent experiments (n = 10–14 mice/group). Bars represent mean ± SEM with individual values shown.

To further characterize the basis for this differential binding, we evaluated antigen expression on human neurons. GD2 was expressed on neurons, albeit at a lower antigen density than HR-NBL cells, whereas B7-H3 expression was virtually absent (**Fig. 5c**). Consistent with these expression patterns, dinutuximab bound to neurons at concentrations sufficient to mediate ADCC *in vitro* (based on studies shown in **Fig. 2**), while INV724 exhibited minimal binding even at the high concentrations tested (**Fig. 5d**).

Using flow cytometry, we also assessed antibody binding to cells isolated from murine dorsal root ganglia. Dinutuximab demonstrated substantial binding to dorsal root ganglion cells, whereas INV724 exhibited only minimal binding above background levels (**Fig. 5e-f, Fig. S4**). Together, these findings indicate that dual-antigen recognition effectively prevents binding of the GD2xB7-H3 bsAb to neural tissues that express GD2 but lack B7-H3.

Because antibody binding to GD2-expressing nerve tissue has been associated with treatment-related pain, we next investigated whether the reduced nerve binding observed with INV724 translated into decreased pain-related behaviors *in vivo*. Many preclinical pain models rely on stimulus-evoked measures, such as heat or mechanical stimulation, which have shown limited translational relevance to clinical pain outcomes.^33, 34^ We therefore employed a model of spontaneous pain, as spontaneous pain measures may more closely reflect the human experience of pain following systemic administration of anti-GD2 antibodies.^35^ The burrowing assay measures species-typical housekeeping behavior in mice that is important for maintaining normal well-being and has been proposed to reflect aspects of quality of life analogous to activities of daily living in humans (**Fig. 5g, Fig. S5**).^36, 37, 38, 39^ Reduced burrowing activity is indicative of stress and discomfort associated with pain. As shown in **Fig. 5h**, mice treated with dinutuximab displaced significantly less bedding than mice treated with rituximab (control mAb) or sham-treated controls. In contrast, burrowing activity in INV724-treated mice did not differ from that of either control group and was significantly different from that seen with dinutuximab. These findings indicate that, unlike dinutuximab, INV724 did not induce measurable pain-related behavioral changes in this model.

## DISCUSSION

By requiring co-expression of two distinct tumor antigens, GD2 and B7-H3, this bispecific antibody strategy enhances tumor specificity beyond that achievable with conventional monospecific antibodies. Although GD2 is highly expressed on NBL, it is also present on peripheral nerves and contributes substantially to the dose-limiting neuropathic toxicity associated with anti-GD2 therapy. In contrast, B7-H3 is broadly expressed on NBL (albeit at lower density than GD2) but is not meaningfully detected on nervous tissue. Here, we demonstrate that GD2xB7-H3 bsAb binds efficiently to cells co-expressing both antigens while exhibiting no substantial binding to neural tissues that express GD2 alone. Importantly, INV724 retains antitumor activity comparable to dinutuximab in both *in vitro* ADCC assays and *in vivo* tumor models, while eliminating detectable binding to rat sympathetic ganglia, human peripheral nerves, and murine dorsal root ganglia. Consistent with these findings, INV724 did not induce pain-associated behavioral changes *in vivo* in treated mice Together, these data suggest that dual-antigen targeting can uncouple antitumor efficacy from a major mechanism of anti-GD2-associated toxicity. Clinical phase I–II studies will be required to determine safety and efficacy of this INV724 bsAb in patients.

The current maintenance regimen for patients receiving dinutuximab involves four consecutive daily 4-20 hour intravenous infusions, followed by four additional 4-day cycles administered every 3-4 weeks if tolerated. Yet, the administration of dinutuximab is associated with a high-degree of toxicity, including grade 3-4 neuropathic pain reported in ∼52% of patients.^1, 2^ In many cases, this pain limits the ability to complete full-dose therapy. Reported symptoms include generalized pain as well as abdominal, extremity, and back pain, along with neuralgia and arthralgia.^1, 40^ Consequently, most patients require concurrent opioid analgesia, and some require additional interventions, such as IV lidocaine or sedation to alleviate the acute painful side-effects.^41^ By improving tumor-specificity and virtually eliminating binding to nervous tissue, INV724 has the potential to substantially reduce these neuropathic side effects. If the reduction in nerve binding observed preclinically translates clinically, INV724 may reduce the need for opioid and intravenous analgesic support, potentially facilitating outpatient administration, improving quality of life during therapy, and increasing treatment tolerability.^42^

Prior efforts to reduce anti-GD2-associated neuropathic pain have included engineering antibody modifications to reduce complement activation. The hu14.18K322A variant incorporates a point mutation in the IgG heavy chain that decreases complement activation and has been evaluated clinically for this purpose. Because complement contributes to anti-GD2-associated pain, this strategy was intended to mitigate toxicity.^6, 9, 10, 43^ Patients receiving hu14.18K322A experience reduced pain compared with dinutuximab, and the maximum tolerated dose is approximately 60 mg/m² per dose (2–3-fold higher than dinutuximab monotherapy). However, neuropathic pain is still observed and contributes to dose limitation.^9, 10, 43^ Other approaches include an IgA-based anti-GD2 antibody designed to promote ADCC (particularly via neutrophils) without complement activation, which has shown reduced pain in preclinical models but has not yet entered clinical testing.^44^ Similarly, a bispecific anti-GD2xCD3 T cell engager lacking complement activation has demonstrated potent preclinical activity but has not been clinically evaluated. Additional work has identified O-acetyl-GD2 as a tumor-associated variant preferentially expressed on NBL but not peripheral nerves, and antibodies targeting this antigen have shown antitumor activity without inducing pain in preclinical studies, although clinical data are lacking.^31, 45, 46^ Collectively, these findings suggest that complement activation contributes to anti-GD2-associated pain but is unlikely to be the sole mechanism.^9^ In contrast, antibodies such as rituximab and cetuximab, which do not bind peripheral nerves, are not associated with neuropathic pain.^12, 30, 47, 48, 49^ The persistence of neuropathic symptoms with complement-reduced antibodies, together with the absence of neuropathic toxicity associated with antibodies that do not bind peripheral nerves, supports an important role for direct nerve targeting in the pathogenesis of anti-GD2-associated neuropathic toxicity. These observations provide a mechanistic rationale for dual-antigen strategies that eliminate monospecific GD2 recognition, preventing binding to nerves while preserving tumor targeting for antitumor efficacy.

An alternative strategy for NBL targeting is the use of antibodies against other, non-GD2, tumor-associated antigens; one example is B7-H3, which is currently under clinical evaluation.^50^ Because antibody efficacy in NBL is largely mediated through ADCC and antibody-dependent cellular phagocytosis, we compared a conventional bivalent anti-B7-H3 antibody (I7-01) with INV724 and anti-GD2 antibodies. We observed greater ADCC activity with INV724 and anti-GD2 antibodies compared with anti-B7-H3 antibody. This may reflect (i) higher surface density of GD2 relative to B7-H3 on most NBL and (ii) spatial differences in antigen accessibility, whereby GD2, as a membrane-proximal glycolipid, may position bound antibody closer to the cell surface than the larger extracellular domain of B7-H3. Consequently, monospecific anti-B7-H3 antibodies may have limited binding on cells with low antigen density. In contrast, once the anti-B7-H3 Fab of INV724 binds its target, the anti-GD2 Fab can rapidly engage abundant membrane-embedded GD2, promoting high-avidity dual engagement and enhanced ADCC. Thus, INV724 appears to achieve a favorable balance between specificity and functional potency. By requiring dual-antigen recognition, it restricts binding to tumor cells while leveraging the high surface density of GD2 to maintain strong avidity and efficient effector-cell engagement.

Recent clinical data indicate that anti-GD2 antibodies may be more effective when combined with chemotherapy.^51^ Based on these findings and other current insights, anti-GD2 antibody has shown more rapid partial and complete responses when added to induction chemotherapy for newly diagnosed NBL patients at St. Jude.^6, 8^ Because of this potential benefit for anti-GD2 antibody during the induction chemotherapy and the efficacy shown for relapsed patients when anti-GD2 is given with chemotherapy, the Children’s Oncology Group is moving to a randomized trial, based on a successful pilot trial, that includes dinutuximab in maintenance and randomizes patients to induction with or without anti-GD2 [NCT06172296].^8, 51, 52^ This expanding clinical use across treatment phases further highlights the potential relevance of INV724 if reduced toxicity enables broader and more intensive administration.

Finally, the reduced neuropathic toxicity observed with INV724 may create opportunities to optimize treatment duration and scheduling rather than simply increasing dose intensity. Current anti-GD2 regimens are frequently associated with significant pain and treatment burden, which can limit completion of therapy and necessitate extensive supportive care. Notably, some of our prior clinical analyses have demonstrated a positive relationship between anti-GD2 antibody exposure and clinical outcome, with higher antibody exposure correlating with improved antitumor responses across multiple trials.^6, 14, 51^ Therefore, if neuropathic toxicity is a primary determinant of dose limitation, the improved tolerability of INV724 could permit greater cumulative exposure than is currently achievable with conventional anti-GD2 antibodies. If the improved tolerability observed preclinically translates clinically, INV724 may enable prolonged treatment courses, extended maintenance strategies, or more flexible administration schedules that increase cumulative therapeutic exposure while maintaining patient quality of life. These possibilities would require prospective evaluation in clinical studies.

Several limitations should be acknowledged. First, the present studies were conducted in preclinical models, and the extent to which reduced nerve binding will translate into reduced neuropathic toxicity in patients remains to be determined. Second, although the spontaneous pain model used here has translational advantages over stimulus-evoked assays, it cannot fully recapitulate the clinical experience of anti-GD2-associated pain. Third, while this dual targeting of GD2 and B7H3 together provides a substantial increase in the tumor specific binding of INV724, the fact that the antigen density of GD2 is roughly 200-fold that of B7-H3 may influence the amount of INV724 that can bind to tumor cells in vivo compared to the amount of dinutuximab that can bind in vivo. Although substantially more dinutuximab can bind NBL cells than INV724 under saturating in vitro conditions, the relevance of this observation to antitumor efficacy in vivo remains unknown because it depends on the concentrations of antibody achieved within the tumor microenvironment. Future studies will evaluate tumor interstitial concentrations of INV724 and dinutuximab in mice, as well as the extent of in vivo tumor binding following clinically relevant dosing, to determine whether these differences influence antitumor activity. In addition, the impact of dual-antigen targeting on long-term pharmacokinetics, biodistribution, and antitumor activity across heterogeneous NBL populations and a broader range of antibody doses will require further investigation. This work is underway and will be the focus of a subsequent report.

Regardless of whether higher doses of INV724 further enhance efficacy as a naked bispecific agent compared to dinutuximab, the marked antitumor specificity of this dual targeted reagent may make it an attractive platform for delivery of therapeutic payloads directly to NBL while minimizing exposure to normal tissues, including nerves. This GD2 x B7-H3 format could be adapted to an antibody-drug-conjugate (ADC) or as a carrier of therapeutic radio-nuclides as radio-pharmaceutical therapy (RPT).^53, 54^ In both settings, localized delivery of cytotoxic payloads or therapeutic radionuclide within the tumor microenvironment may provide a “bystander effect” capable of eliminating neighboring tumor cells that have reduced or lost expression of GD2, B7-H3, or both antigens.^55^ This might prevent selection of antigen-loss variants that escape ADCC mediated therapy using a naked antibody like INV724 or dinutuximab.

In summary, these preclinical studies support further evaluation of the GD2×B7-H3 bispecific antibody INV724. By requiring co-expression of two tumor-associated antigens, this approach preserves the potent antitumor activity of GD2-directed immunotherapy while eliminating detectable binding to peripheral nerve tissues and reducing pain-associated behavioral changes *in vivo*. Beyond NBL, these findings illustrate the potential of dual-antigen targeting strategies to improve the therapeutic index of antibody-based cancer immunotherapy by enhancing discrimination between tumor and normal tissues.^56^ Such tumor-selective recognition may also provide a versatile platform for future targeted delivery approaches, including antibody-drug conjugates, radiopharmaceuticals, and other payload-bearing therapeutics.

## METHODS

### Construction of B-Body Bispecific Antibodies

Bispecific antibodies (bsAbs) were created from GD2 and B7-H3 and GD2 antibodies as depicted in **Fig. 1a** by reformatting the B7-H3 antibodies into the first and second polypeptide chains and GD2 into the third and fourth polypeptide chains in an antibody construct. The four polypeptide chains were transiently transfected into HEK cells to produce the antibodies. The bispecific antibodies secreted into the cell culture medium were subsequently isolated with an anti-CH1 affinity capture resin followed by polishing using a strong cation exchange resin. Bispecific constructs in which either the B7-H3 or the GD2 arm was replaced with a non-binding arms were also created.

### Production of INV721 and INV724 bsAbs

INV721 and INV724 bsAbs (Invenra) were produced from CHO cell lines stably expressing the respective gene sequences. For the afucosylated version (INV724), the expression of alpha-(1,6)-fucosyltransferase was inhibited.

### Cell Lines

All cell lines used were routinely monitored for mycoplasma by PCR testing as previously described^57^, and complete culture medium was made by supplementing RPMI-1640 or DMEM with 10% heat-inactivated FBS, 2 mmol/L l-glutamine, 100 U/mL penicillin, and 100 µg/mL streptomycin.

Human NBL cell lines NGP, NB69, NB-EBc1, IMR5, NLF, were kindly provided by Dr. John Maris of the Children’s Hospital of Philadelphia, and L-A-N-1, L-A-N-5 were provided by Dr. Patrick Reynolds of Texas Tech University. Human osteosarcoma cell line, Saos-2, was provided by Dr. Mario Otto of Phoenix Children’s Hospital. LA-N-1, LA-N-5, NGP, NB69, NB-EBc1, IMR5, NLF, SK-NA-S and Saos-2 cell lines were grown in RPMI 1640 and CHLA-20 in DMEM.

M14 melanoma cells (RRID: CVCL_1405) were obtained from Dr. Ralph Reisfeld (Scripps Research Institute). Although this cell line was previously reported as M21 in publications from our laboratory, recent cell line authentication identified it as M14. Human melanoma cell lines (MRA_Mel3, MRA_Mel4, MRA_Mel7 and MRA_Mel13) were kindly provided by Dr. Mark Albertini of the University of Wisconsin-Madison, were grown in RPMI 1640. A673 cells were purchased from ATCC and grown in DMEM.

The murine NXS2 neuroblastoma cell line (kindly shared by Ralph Reisfeld, PhD, The Scripps Research Institute, La Jolla, CA, and then maintained by Alice Yu, MD, University of California, San Diego, CA) is a moderately immunogenic, highly metastatic, GD2-positive line.^58^ NXS2 is a hybrid between GD2-negative C1300 (a neuroblastoma tumor that spontaneously arose in A/J mice^59^) and GD2-positive murine dorsal root ganglion cells (C57Bl/6 J background), but does not express C57Bl/6 H-2 and therefore grows in immunocompetent A/J mice. NXS2 cells were grown in DMEM medium.

The murine B78-D14 (“B78”) melanoma cell line is a poorly immunogenic line derived from B78-H1 originating from the B16 line, and was gratefully obtained from Ralph Reisfeld, PhD at The Scripps Research Institute in La Jolla, CA.^60, 61, 62^ B78-D14 cells have functional GD2/GD3 synthase and express the disialoganglioside GD2.^60, 61^ B78 cells were grown in RPMI 1640, and periodic treatment with hygromycin B (50 μg/mL) and G418 (400 μg/mL) was used to maintain GD2 expression.

### Cell Line Modifications

B78s were modified to knock-out expression of murine B7-H3 via CRISPR. To do this, B78s at 50% confluency in a 96 well plate were treated with single guide RNA (sgRNA) targeting B7-H3 (TrueGuide™ Synthetic; ThermoFisher, cat. no. CRISPR142889_SGM) using TrueCut^™^ Cas9 Protein v2 (ThermoFisher, cat. no. A36496) and Lipofectamine^™^ CRISPRMAX^™^ Cas9 Transfection reagent (ThermoFisher, cat. no. CMAX00001) following the manufacturer’s protocol. After single-cell cloning, expression of mouse B7-H3 (mB7-H3) was assessed by flow cytometry compared to the original untransfected control B78 cells. For flow cytometry, cells were labeled with anti-mouse B7-H3-APC antibody (Biolegend, cat. no. 135608) for 30 min, washed and analyzed on an Attune NxT Flow Cytometer (ThermoFisher). B78 clones that showed at least 50% decrease in mB7-H3 were treated again with B7-H3 sgRNA, followed by single-cell cloning and assessment by flow cytometry for mB7-H3 expression. A B78 clone that had lost expression of mB7-H3 (B78^mB7-H3-^) was selected following two treatments with B7-H3 sgRNA treatment.

After confirmation by flow cytometry that B78^mB7-H3-^cells did not express mB7-H3, human B7-H3 (hB7-H3) was transduced into B78^mB7-H3-^ and NXS2 (that still had mB7-H3) by lentiviral transduction using pLV-mCherry:T2A:Puro-EFS>hCD276 [NM_001024736.1] (VectorBuilder; Vector ID VB170825-1084zey). Successfully transduced cells were selected for using puromycin (4 µg/mL for B78, 2 µg/mL for NXS2). For B78s, stably transduced cells, referred to as B78^mB7-H3-/hB7-H3+^, were then single-cell cloned and tested for GD2 and hB7-H3 expression. Five mB7-H3-negative B78 clonal lines were established based on variable expression of hB7-H3 and GD2 and used for different purposes. Clone (Cl.) 6 (hB7-H3+/GD2+) was used for *in vivo* efficacy testing. Others were used for *in vitro* specificity, *in vitro* efficacy/ADCC and PET imaging studies, including Cl. 8 (hB7-H3-/GD2-), Cl. 13 (hB7-H3+/GD2+), Cl. 14 (hB7-H3+/GD2-), Cl. 25 (hB7-H3-/GD2+). For NXS2, stably transduced NXS2+hB7-H3 cells were sorted, single-cell cloned and tested for GD2 and human B7-H3 expression.

For *in vivo* IVIS imaging, CHLA-20 and SK-NA-S cells were transduced as above with lentiviral transduction using pLV-Puro-EFS>TurboGFP:IRES:Luciferase (VectorBuilder; Vector ID VB180725-1103sza). Successfully transduced cells were selected with puromycin (2 µg/mL).

### Cell Binding

The GD2xB7-H3 bsAb INV721 or it afucosylated version, INV724, and the corresponding GD2xNon-Binding or B7-H3xNon-Binding bsAbs were tested by flow cytometry for binding to B78 murine melanoma cell lines overexpressing GD2 +/-hB7-H3 or to various human NBL, melanoma or osteogenic sarcoma cell line cells. Cells were incubated for 60 minutes at 4°C with a dilution series of the bsAbs ranging from 333 nM to 0.02 nM prepared in PBS (Corning, #21-031-CV). They were then washed with PBS and resuspended in a secondary antibody solution of AF488 goat anti-human IgG Fab Fragment (Jackson Immuno Research, #109-547-003) diluted in PBS. After a 30-minute incubation at 4°C, the cells were washed, resuspended in cold PBS, and mean fluorescence intensity (MFI) was measured via flow cytometry.

### Measurement of Binding Affinities

Monovalent affinities to GD2 and B7-H3 were determined by biolayer interferometry on the OctetQK384 system (Pall ForteBio). Biotinylated GD2 or B7-H3 was immobilized on streptavidin sensors. The antigen-immobilized sensor was submerged into a solution containing various concentrations of bsAb ranging from 150-10,000 nM for GD2 or 3-200 nM for B7-H3. The real-time association and dissociation curves of the bsAb molecule were fit using ForteBio software in a 1:1 binding model with global or local fit to obtain association (kon) and dissociation (koff) rates. *K*d was calculated from these rate constants.

### Antibody Dependent Cellular Cytotoxicity (ADCC)

Target tumor cell lines transduced to express nuclear localized mKate2 (NLS-mKate2) were plated in 20 µL/well in a 384-well round bottom plate (Corning, #4516) at 100 or 250 cells/well (depending on cell size). Cells were incubated at 37°C for 48 hours to allow spheroids to form. During that incubation, peripheral blood mononuclear cells (PBMCs) were isolated from healthy donors as previously described and incubated overnight at 37°C in RPMI 1640 media supplemented with IL-2 (200 U/mL).^3^ PBMCs were centrifuged at 350g for 10 minutes, washed with 50 mL of PBS to remove IL-2, and resuspended in RPMI 1640 without IL-2. 2,500 PBMCs were added to each well (each spheroid) in the 384-well plate in 20 µL of media and antibodies were added in 10 µL/well. Staurosporine was included as a positive (maximum) tumor death control at a 1 mM/well. The final volume of all wells was brought to 50 µL with media and the plates were placed in an IncuCyte S3 system (Sartorius) and imaged every 4 hours using the Spheroid Module.

### FcγR Genotyping Assays

Genotyping for Fc*γ*R3A, Fc*γ*R2A and Fc*γ*R2C and categorical determinations for high-affinity Fc*γ*Rs vs low-affinity Fc*γ*Rs were performed as previously described.^63, 64, 65^

### Antibody Internalization Assays

Antibodies were labeled with pHrodo^TM^ iFL Red (ThermoFisher) per the manufacturer’s protocol. Antibodies were plated together with NBL cell lines and imaged every 3-6 hours for 48 hours via an IncuCyte S3 live cell imaging platform.

### Animal Models

Female 6-8-week-old C57BL/6 (Taconic Farms, strain B6NTac) and A/J mice (Jackson Labs; Strain#: 000646) were used for these studies involving syngeneic tumor models (B78 and NXS2, respectively). Female 6-8-week-old nude mice (Jackson Labs, Strain#: 007850; RRID:IMSR_JAX:007850) were used for studies involving human NBL tumors (CHLA-20). Female 6-8-week-old NOD.Cg-Rag1^tm1Mom^ Il2rg^tm1Wjl^/SzJ mice (NRG; Jackson Labs, Strain#: 007799) were used for PET imaging studies of NGP neuroblastoma tumors (described below). Mice were housed in the University of Wisconsin-Madison animal facilities at the Wisconsin Institutes for Medical Research. Mice were used in accordance with the *Guide for Care and Use of Laboratory Animals* (NIH publication 86-23, National Institutes of Health, Bethesda, MD, 1985).

Intradermal tumors were established by injecting 2x10^6^ B78 Cl.6, or NXS2+hB7-H3 in 0.1 mL of Matrigel (Sigma, # 354248) with PBS into the shaved flank of mice.^66^ Tumor diameters were measured, and tumor volume (mm^3^) was calculated as: ^1^ (*tumor lengt*ℎ × *tumor widt*ℎ)^2^. One day before radiation therapy, mice were randomized according to tumor volumes and assigned to experimental groups.

Intravenous (IV) metastatic tumors were established by injecting 5x10^5^ luciferase-expressing CHLA-20 or SK-NA-S cells in 0.1 mL of PBS into the tail vein of mice. Tumor development was monitored weekly by bioluminescent imaging with Perkin Elmer IVIS Spectrum In Vivo Imaging System and Living Image Software (Caliper Life Sciences, Hopkinton, MA) was used for image processing. Bioluminescence was measured 15 minutes following intraperitoneal injection of 200 µL (150 mg/kg) of luciferin.

### Burrowing Method to Monitor Pain

A mouse burrowing assay was used as a correlative metric to assess pain based on previously published methods.^36, 38, 39, 67^ Mouse burrowing apparatuses were constructed in house by modifying a standard red mouse tunnel (VWR, cat no. Bio-Serv K322) to seal off one end with a plastic cap (**Fig. S5a**). On Day 0, 4 cages of mice (labeled A-D) were given 4 hours to acclimate to an empty burrowing apparatus. Burrowing apparatuses were labeled according to cage (A-D) and saved for testing on subsequent days.

On each subsequent test day, 4 test cages were prepared with labels (A-D) and separated by noise/visual barriers within a biosafety cabinet with the blower turned on **(Fig. 5g, Fig. S5b)**. Burrowing apparatuses were filled with 30g of bedding (5g from home cage and 25g fresh bedding) and placed in their corresponding cages (**Fig. 5g**). One mouse at a time from each cage was moved from its home cage and placed into the corresponding test cage for 15 minutes. Following each test, mice were returned to their home cage and the burrowing apparatus was weighed to measure the amount of bedding removed (difference from initial 30g). All burrowing assessments were performed blinded, and bedding was stored by cage in plastic bags for the remaining days of the study.

On Day 1 at 9am, mice were tested for pre-burrowing to ensure they would burrow. Mice were left alone for 15 min, and the amount of bedding removed from the burrow was measured. Mice were then returned to their home cage. On Day 2 at 9am, mice were tested for pre-treatment burrowing at time 0 as described above. Mice that removed >20g of bedding on Day 2 were enrolled in the study.

Following the screen on day 2, enrolled mice were injected IV (tail vein) with 200 µL of treatment: dinutuximab (250 µg), INV724 (250 µg), rituximab (250 µg), or a needle-poke sham control. All cages included at least one mouse from each treatment group to control for emotional contagion. One hour after treatment administration, burrowing activity was tested, and the percent change in activity (by bedding weight) from the Day 2 pre-screen (before treatment) to 1 hour post-treatment was calculated. Burrowing studies were repeated 3 separate times, with 3-5 mice enrolled per treatment group during each study for a final total of 10-14 mice/group.

### PET Imaging of Tumors

NRG mice were injected with 10^6^ NGP tumor cells intradermally^66^ on the lower right flank. Separately, C57Bl/6 mice were injected with 10^6^ B78^mB7-H3-^cells intradermally^66^: B78^GD2+/B7-H3-^(Cl. 25) on the lower left flank or B78^GD2+/B7-H3+^ (Cl. 13) on the lower right flank (**Fig. 4g**). When the tumors reached ∼100-300 mm^3^, the mice were injected with ^89^Zr-labeled antibodies prepared as described previously.^68^ Briefly, p-SCN-Bn-Deferoxamine (Df) was conjugated by thiourea linkage to the antibody at pH ∼8.5 and purified using size exclusion chromatography. Conjugated antibody was radiolabeled with ^89^Zr produced on the UW-Madison GE PETtrace cyclotron by using the ^89^Y(p,n)^89^Zr reaction. For radiolabeling, ^89^Zr-oxalate was added to the chelator conjugated antibody and incubated in 1 M HEPES buffer (pH=7.5) at 37°C for 1 hour at a ratio of 100 μg antibody per mCi of ^89^Zr. Radiolabeled antibody (^89^Zr-Df-INV721) was purified using size exclusion chromatography. Mice were injected IV with 5.55-9.25 MBq (150-250 μCi) for PET imaging studies. PET/CT images were decay corrected and images are normalized to percent injected dose per gram of tissue (%ID/g) using the Inveon Research Workspace by manually drawing volumes-of-interest. Additional imaging, avidity, selection and biodistribution studies were performed using other ^89^Zr-labeled GD2xB7-H3 bispecific antibodies.^69^

### In Vivo Tumor Model Efficacy Studies

Mice bearing GD2+/B7-H3+-expressing melanoma (B78^GD2+/B7-H3+^, Cl. 6) or neuroblastoma (NXS2) tumors were treated with our *in situ* vaccine regimen: external beam radiation therapy (12Gy, Day 1) followed by intratumoral injection of IL-2 (75K U/dose) with or without INV724 or dinutuximab (40µg/dose, Day 5-9). Mice were monitored for tumor growth via twice weekly caliper measurements. Mice were euthanized when either the length or the width of a tumor reached 20 mm and tracked for survival.

Nude mice were injected IV with CHLA-20-luciferase tumors on Day 0, followed by retro-orbital injection of INV724 or dinutuximab (25 µg/dose) in 0.1 mL of PBS, or PBS alone, on Days 1, 4, 7 and 10. Retro-orbital injections were performed using sterile technique under isoflurane anesthesia.

### Antibody Titrations

Tumor cells were harvested and resuspended at 5x10^6^ cells/mL in flow buffer. From this suspension, 50 µL of cells were added to flow cytometry tubes, for a total of 250,000 cells/tube. Serial dilutions of primary antibody (INV724 or dinutuximab) were created, and 50 µL of each antibody dilution was added its respective tube. Cells were incubated for 30 minutes in the dark at 4°C and washed with flow buffer (PBS + 2% FBS). A secondary master mix was made by diluting anti-Fab-PE (109-117-008; Jackson ImmunoResearch labs) 1:50 in flow buffer, and 100 µL of the master mix was added to each tube and incubated for 20-30 minutes in the dark at 4°C. Samples were washed with flow buffer, DAPI was added, and cells were subjected to flow cytometry on an Attune NxT (ThermoFisher) or an IntelliCyt iQue Screener PLUS (Sartorius) to determine the mean fluorescence intensity (MFI).

### Antigen Density Testing

Cells were harvested and added to flow tubes (200,000/100 µL flow buffer) for staining with GD2-APC (BioLegend, 357306) or CD276 APC (BioLegend, 351006). Quantum Simply Cellular anti-mouse IgG beads (Bangs Laboratories, 815A) were prepared in flow buffer in microcentrifuge tubes and stained at the same time. Amounts of antibody added to the beads and to each cell line were determined previously by antibody saturation titration (additional 50% antibody volume added until less than a 10% increase in fluorescence is obtained=antibody saturation). Cells and beads were incubated for 30 minutes in the dark at 4°C and washed with flow buffer. Beads were washed an additional two times as per manufacturer’s recommendations. DAPI was added to cell samples before running on an Attune NxT flow cytometer and beads were collected after the cells at the same instrument settings. Bangs QuickCal template for BD Relative Linear Channels (1-10,000 channels, for Log Data) and bar (range) gates for the bead populations (about 1000 beads per tube collected) were used for Antigen Binding Capacity (ABC) analysis and the unstained control ABC result was subtracted from the test ABC result for each cell line. If monovalent antibody to receptor binding is presumed, the ABC value is the number of surface receptors.

### Human Neuron Titrations and Antigen Density Testing

For human neuron titrations, human neurons (ScienCell, 1520-10) were grown in Neuronal Medium (ScienCell; Cat. No. 1521) on poly-L-Lysine coated plates according to the manufacturers protocol. Neurons were thawed according to ScienCell Research Laboratories’ protocol, and titrations and antigen density testing were immediately performed as above.

### Immunofluorescent Staining of Tissue

Rat peripheral nerve tissue and CHLA-20 tumors grown in nude mice were harvested, embedded in OCT without fixation, and frozen on liquid nitrogen. 10 µm cryosections from the embedded tissues were collected onto Colorview adhesion slides (StatLab). OCT-embedded human peripheral nerve tissue (specimen 407E) was a kind gift from Ambsio and sectioned at 7 µm per tissue onto Superfrost slides (ThermoFisher). Following a slow thaw after removal from -80°C storage, slides were incubated in cold acetone (maintained at -20°C) for 10 minutes and then dried for 10 minutes at room temperature (RT). Slides were rinsed under water for 10 minutes, rinsed with PBS 3 times, and blocked with 10% FBS for 60 minutes at RT. For rat nerve tissue, primary INV724 (Invenra) or dinutuximab antibodies (commercial grade, Unitixun) at 3 µg/mL in 1% FBS, or unstained sections in 1% FBS block, and incubated overnight at 4°C. Slides were washed 3 times in PBS, and 1% FBS with 1 µg/mL of secondary anti-human-IgG-AF555 (A-21433, LifeTech) was added and incubated for 60 minutes at RT. For human peripheral nerve staining, sections were stained with primary INV724+V5 tag or dinutuximab+V5 tag antibodies (Invenra) at 3 µg/mL in 1% FBS, or unstained sections in 1% FBS block, were incubated overnight at 4°C. Slides were washed 3 times in PBS, and 1% FBS with 1 µg/mL of secondary anti-V5-AF55 (49355S, Cell Signaling) was added and incubated for 60 minutes at RT. Slides were washed 3 times with PBS, and fixed in 10% neutral buffered formalin for 10 minutes. Slides were washed 3 times in PBS, and incubated with phalloidin-AF488 (15 nM, A12379, ThermoFisher) for 30 minutes at RT. Slides were washed with PBS 3 times, with a drop of DAPI being added with the third wash. Slides were cover slipped with Prolong Gold mounting media (ThermoFisher) and imaged on a confocal microscope (LSM 710). These staining procedures were repeated twice with similar results.

### Dorsal Root Ganglion Flow

Dorsal root ganglion were isolated from 4 naïve C57Bl/6 mice and digested into single cell suspensions.^70^ Single cell suspensions were stained with 1 µg of dinutuximab or INV724 and incubated at RT for 20 minutes. Cells were washed with flow buffer, centrifuged (350g X 5 minutes), and incubated with secondary anti-human IgG1-PE (MA1-10389, ThermoFisher) for 20 minutes at RT. Cells were washed with flow buffer, centrifuged, and incubated with A2B5-AF647 antibody (BioLegend, catalog #150704) to stain dorsal root ganglion nerve cells) for 20 minutes. Cells were washed, centrifuged, and DAPI was added prior to flow cytometry analysis on an Attune NXT (ThermoFisher). Data were analyzed using FlowJo software by gating on live, A2B5+ nerve cells, and assessed for the amount of bound secondary PE antibody by Median Fluorescence Intensity (MFI) to either dinutuximab or INV724. This study was repeated twice with similar results.

### Statistics

Tumor spheroid shrinkage data were compared by one-way ANOVA followed by Tukey’s Post Hoc test for multiple comparisons. For statistical analysis of tumor growth curves, the time-weighted average (area under the volume-time curve, calculated using trapezoidal method) was calculated for each mouse tumor. Time-weighted averages were compared between treatment groups overall by a Kruskal-Wallis test, and since significance was reached, then pairwise by Mann-Whitney-Wilcoxon tests. Survival curves were compared with pairwise log rank tests and response was evaluated by two-sample tests of proportions. Chi-square test was used to assess differences in response rate (i.e., tumor free vs. tumor bearing). Significance was assessed at the alpha = 0.05 level and no adjustments were made to account for inflated type 1 error rate. Analysis was conducted using R version 4.3.1 (2023-06-16).

## Data Availability Statement

The data generated in this study are available upon request from the corresponding author.

## Acknowledgements

This work was supported by Invenra Inc, The Wisconsin Alumni Research Foundation, Midwest Athletes Against Childhood Cancer, the University of Wisconsin Carbone Cancer Center and research grants from the Pablove Foundation, the HESI-thrive Foundation, the Hyundai Hope on Wheels Foundation, the End Kids Cancer Foundation, The St. Baldrick’s Foundation, The Band of Parents Foundation, The Cure Childhood Cancer Foundation, The Super Jake Foundation, and by public health service grants R35-CA197078, and P01 CA250972 from the National Cancer Institute.

## Author contributions

AKE, PMS, ASF conceived the study and designed the experiments. AKE, ASF, SNR, AH, ZTR, YG, MF, CH, MH, EF, MG, DS, NT, and VS performed the experiments and collected the data. AKE, ASF, SNR, ZTR, YG, MF, and VS analyzed the data. AKE, ASF, SNR, RH, BH, JHD, RG, ALR, JAH and PMS wrote the manuscript. AKE, RH and PMS supervised the project and secured funding. JW, DJG, MB, BW, JWE, and EAS synthesized the chemical compounds. JZ performed the statistical and computational analyses. All authors reviewed and edited the manuscript.

## Inclusion and Ethics Statement

All of the coauthors on this study agree that the research conducted is relevant, and they have fulfilled the criteria for authorship required by Nature Portfolio journals, as their participation was essential for the design and implementation of the study. The roles and responsibilities were agreed among collaborators ahead of the research. All of the research conducted was approved under approved Biological Safety, University of Wisconsin Institutional Review Board and Institutional Care and Animal Use Committee for the University of Wisconsin, Madison. This research was not severely restricted or prohibited in the setting of the researchers, and does not result in stigmatization, incrimination, discrimination or personal risk to participants. Local and regional research relevant to our study was taken into account in citations.

## SUPPLEMENTAL MATERIALS

**Supplementary Figure 1:**
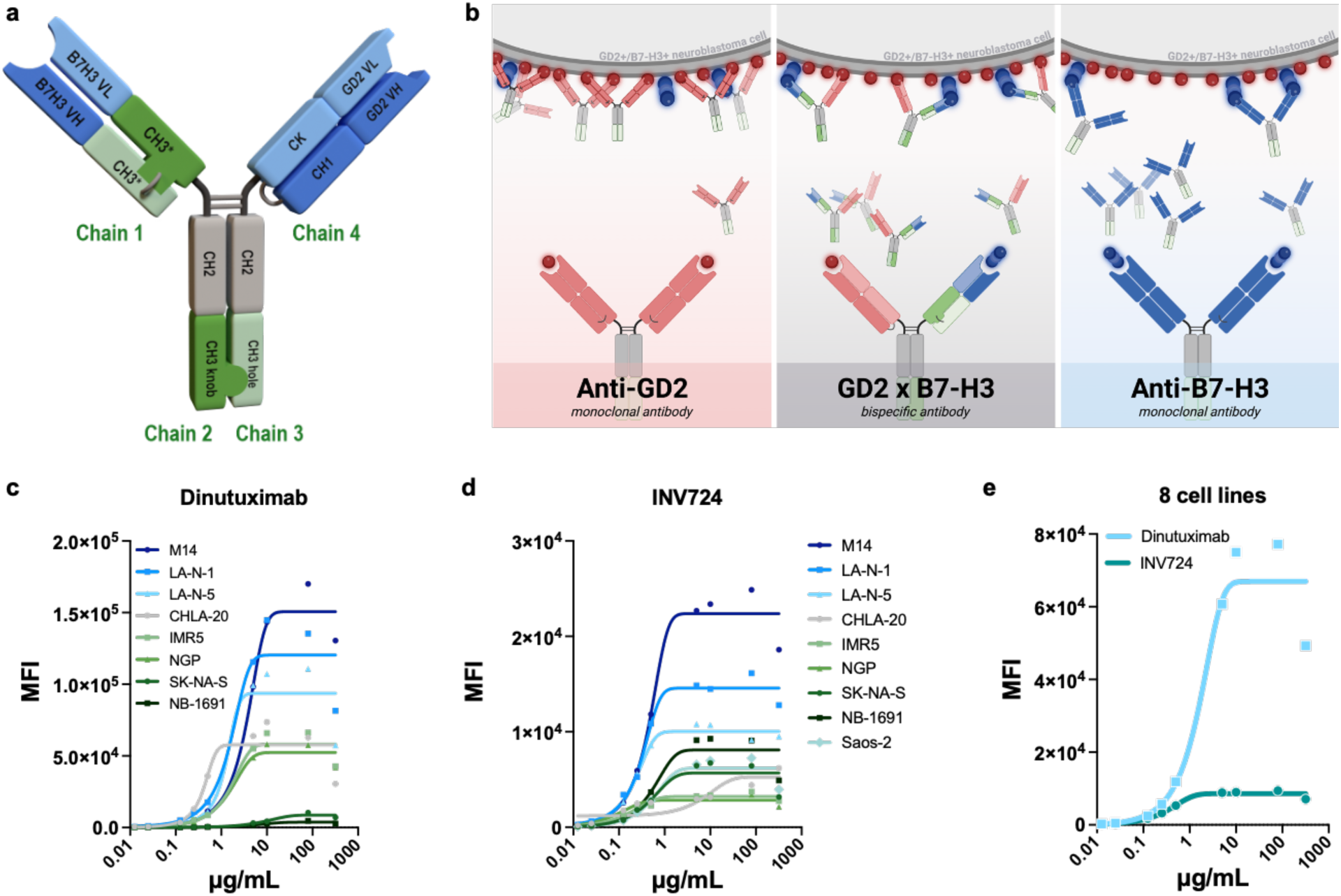
Relative binding of INV724 and dinutuximab to human NBL cells. **a)** Schematic of the 4 immunoglobulin chains used to create INV721 and its afucosylated variant INV724. **b-c)** Flow cytometric analysis of antibody binding to human tumor cell lines. MFI of dinutuximab binding to 8 human tumor cell lines **(b)** and INV724 binding to 9 human tumor cell lines **(c).** Note that the y-axes are displayed on different scales. **d)** Comparison of the mean MFI values for the tumor cell lines shown in B and C, with dinutuximab and INV724 binding displayed on the same scale. **e)** Schematic illustrating the enhanced tumor binding of the B7-H3×GD2 bispecific antibody INV724. GD2 is expressed at high density on NBL cells and can be effectively targeted by anti-GD2 monoclonal antibodies (left), resulting in high-avidity binding and substantial tumor-cell occupancy. In contrast, B7-H3 is often expressed at lower antigen density, limiting the binding and retention of anti-B7-H3 monoclonal antibodies and reducing their capacity to engage immune effector functions (right). By simultaneously targeting GD2 and B7-H3, the bispecific antibody INV724 achieves preferential tumor targeting and enhanced avidity through dual-antigen engagement, resulting in greater tumor-cell binding than an anti-B7-H3 monoclonal antibody alone (center).

**Supplementary Figure 2:**
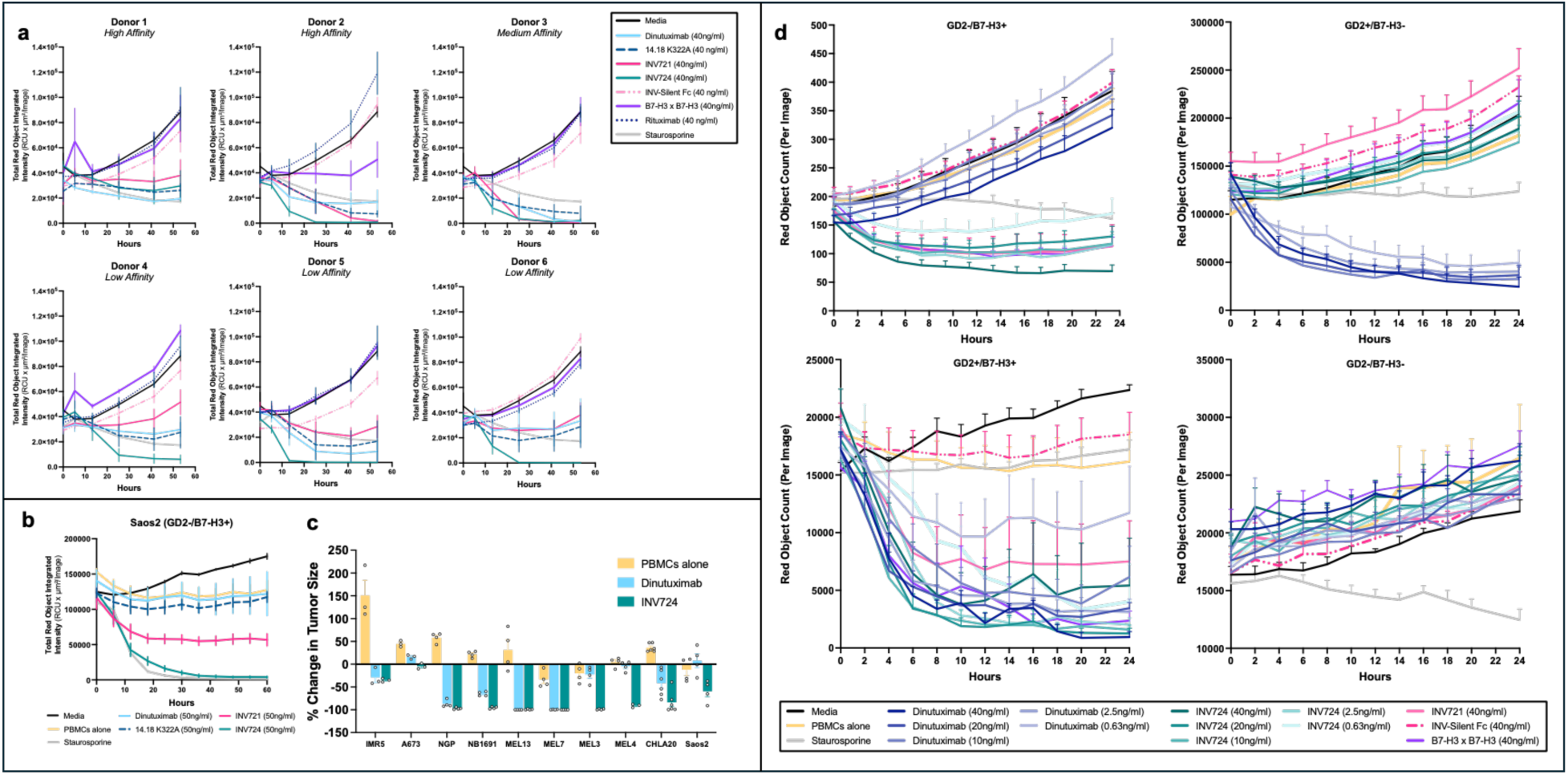
Fc*γ*R genotypes influence ADCC activity. **a)** ADCC assays (via IncuCyte) were performed using PBMCs from six healthy donors and 40 ng/mL antibody. Donors with medium- or high-affinity FcγR genotypes (top panels) demonstrated comparable ADCC activity with INV721 and INV724 regardless of fucosylation status. In contrast, donors with low-affinity FcγR genotypes (bottom panels) exhibited the greatest ADCC activity with the afucosylated antibody INV724. The FcγR3A (V/F) and FcγR2C (C/T) genotypes previously associated with receptor affinity are indicated for each donor.27,28,61,62 Growth curves represent mean ± SEM of four replicate wells. **b)** Percent tumor cell killing at 24 hours from the ADCC assays shown in Fig. 2f. Using a single donor and 50 ng/mL antibody, INV724 induced significantly greater ADCC than dinutuximab across 10 tumor cell lines (Fig. 2f), including four melanomas (MRA-Mel3, MRA-Mel4, MRA-Mel7, and MRA-Mel13), four NBL (NGP, IMR5, NB1691, and CHLA-20), one Ewing sarcoma (A673), and one osteosarcoma (Saos2). **c)** Antibody titration studies (0.63–40 ng/mL) comparing dinutuximab and INV724 in ADCC assays against the B78 GD2/B7-H3 variants shown in Fig. 2h as well as 40 ng/ml of other mAbs.

**Supplementary Figure 3:**
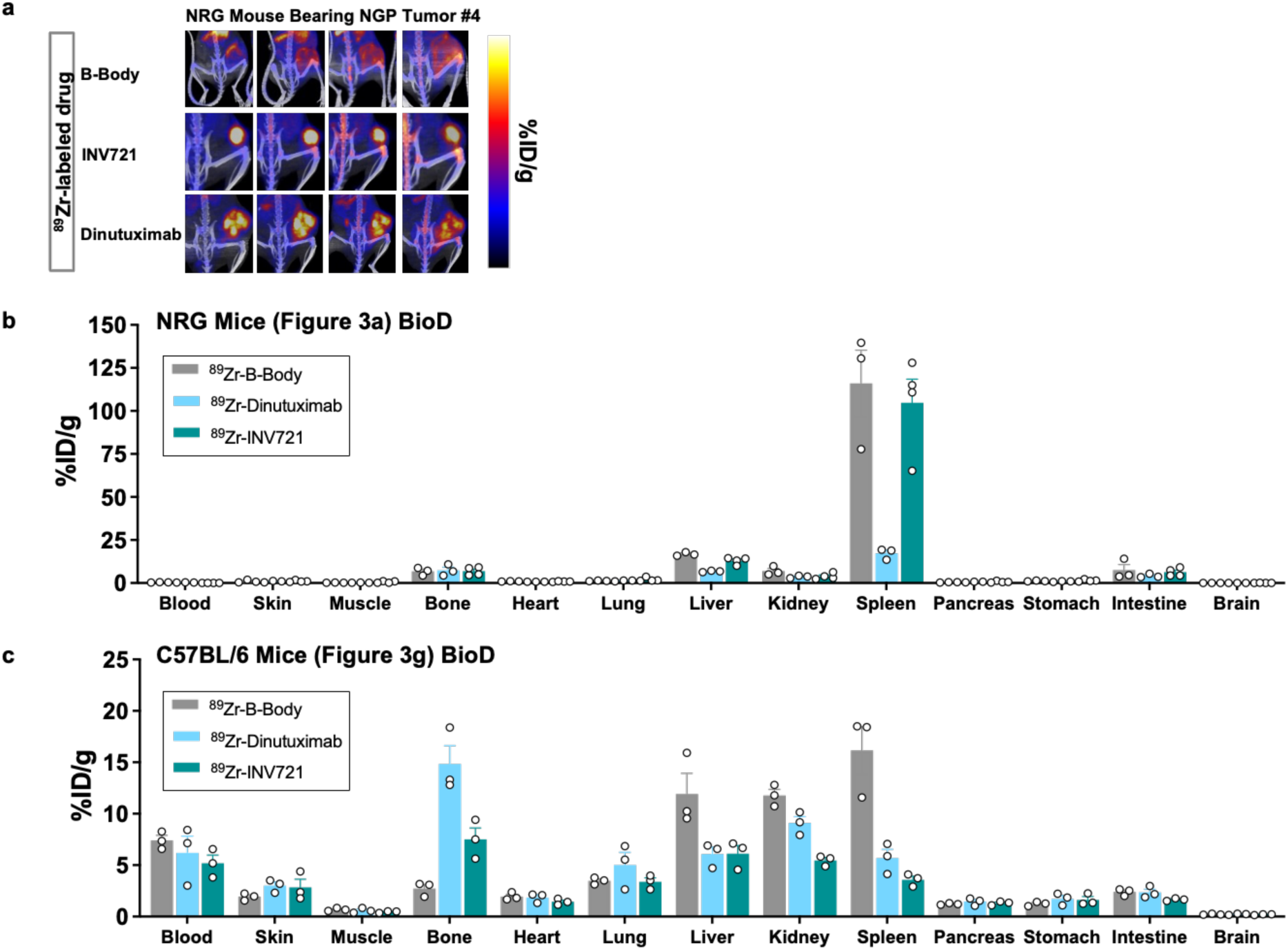
PET Imaging Study Data. **a)** For Fig. 3a-b, four mice per group were analyzed by PET imaging. Three mice per group are shown in Fig. 3A; the fourth mouse from each group is shown here. **b)** In NRG mice bearing NGP tumors, ex vivo biodistribution (BioD) analysis after 72 hrs showed increased splenic accumulation in mice injected with ^89^Zr-INV721 and ^89^Zr-B-Body control antibody, whereas splenic accumulation was not detected in mice administered ^89^Zr-dinutuximab. Nonspecific splenic uptake of radiolabeled human antibodies has been reported in immunodeficient mouse strains and may result from interactions between antibody Fc regions and Fc receptors on innate immune cells.^27^ Given that dinutuximab rapidly internalizes following GD2 binding, whereas INV721 is non-internalizing and the B-Body control antibody lacks antigen-specific binding, the observed splenic accumulation likely reflects nonspecific antibody clearance and retention of the residualizing ^89^Zr radiolabel rather than antigen-specific binding. Data are presented as mean ± SEM. **c)** In C57BL/6 mice bearing B78 flank tumors, BioD analysis of radiolabeled antibody accumulation in tissues.

**Supplementary Figure 4:**
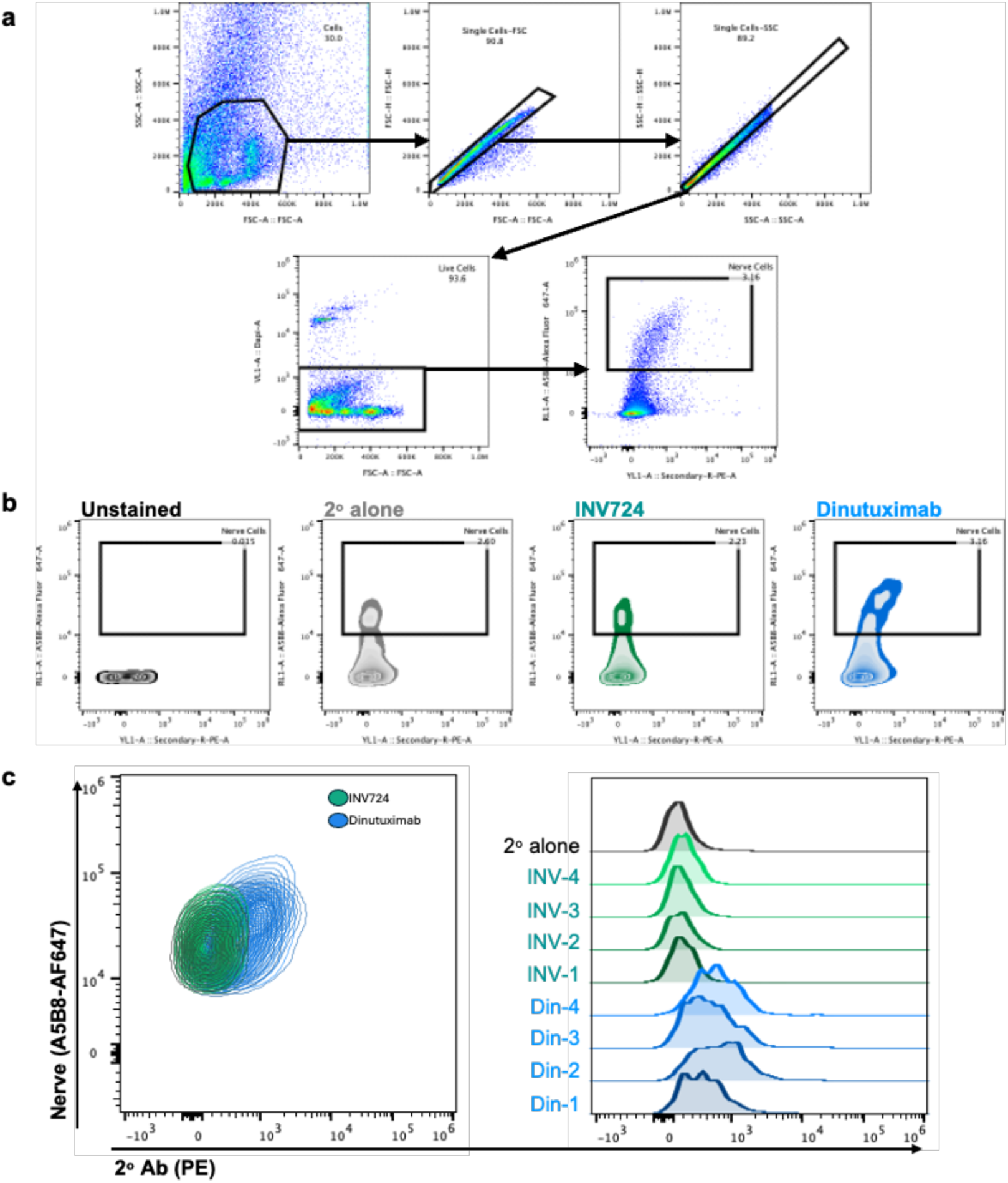
Gating strategy for dorsal root ganglion nerves. **a-c)** Gating strategy for dorsal root ganglion analysis by flow cytometry: **a) (1)** Murine dorsal root ganglion cells were gated on by Forward Scatter Area (FSA) and by Side Scatter Area (SSA). Single cells were then gated on by selecting gates within FSA x FS-Height (FSH) (**2**) and then by SSA x SS-Height (SSH) (**3**), followed by gating on live DAPI-negative cells (**4**). The left plot of A2B5-AF647 Ab x Secondary PE Ab (**5**) shows that based on the no antibody control and the secondary PE Ab only control (no nerve A2B5-AF647 antibody) samples, gates for nerve positive samples were set. **b)** Cells within this gate labeled “Nerve Cells” treated with secondary antibody alone, or INV724 + secondary antibody, or DIN + secondary antibody were used to create the histogram (**c**) and cell contour plots displayed within Fig. 5e**-f**. The results are shown from 4 mice per group (individual mice labeled as DIN 1-4 or INV 1-4).

**Supplementary Figure 5.**
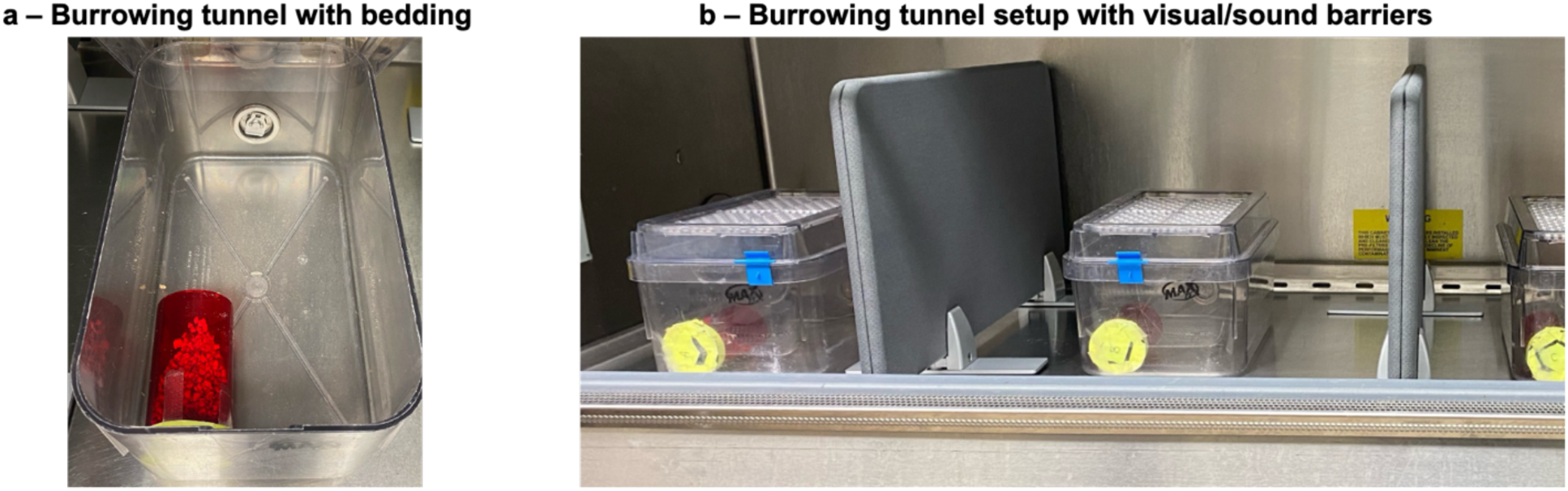
Images of Mouse Burrowing Apparatus. **a**) Burrowing tube containing 30g bedding placed within mouse cage. **b**) Individual burrowing apparatuses housed within cages separated by visual and sound barriers inside a biosafety cabinet to minimize external stimuli during behavioral testing.

